# A 117 year retrospective analysis of Pennsylvania tick community dynamics

**DOI:** 10.1101/433664

**Authors:** Damie Pak, Steven B. Jacobs, Joyce M. Sakamoto

## Abstract

**Background:** Tick-borne diseases have been increasing at the local, national, and global levels. Researchers studying ticks and tick-borne disease need a thorough knowledge of the pathogens, vectors, and epidemiology of disease spread. Three surveillance approaches are commonly used to provide insight into tick-borne disease risk: human disease case surveillance, active tick surveillance, and passive tick surveillance. Long-term passive surveillance can provide up-to-date data on the spatial variability and temporal dynamics of ectoparasite communities and shed light into the ecology of rarer tick species. We present a retrospective analysis on compiled data of ticks from Pennsylvania over the last 117 years.

**Methods:** We compiled data from ticks collected during tick surveillance research, and from citizen-based submissions to the Penn State University Department of Entomology (PSUEnt). Specimens were deposited at the PSUEnt arthropod collections that eventually became The Frost Entomological Museum. While most of the specimens were submitted by the public, a subset of the data were collected through active methods (flagging or dragging, or removal of ticks from wildlife). We analyzed all data from 1900-2017 for tick community composition, host associations, and spatio-temporal dynamics.

**Results:** In total there were 4,491 submission lots consisting of 7,132 tick specimens. Twenty-four different species were identified, with the large proportion of submissions represented by five tick species. We observed a shift in tick community composition in which the dominant species of tick (*Ixodes cookei*) was overtaken in abundance by *Dermacentor variabilis* in the early 1990s, and then replaced in abundance by *I. scapularis*. We analyzed host data and identified overlaps in host range amongst tick species, suggesting potential hubs of pathogen transfer between different tick vectors and their reservoir hosts.

**Conclusions:** We highlight the importance of long-term passive tick surveillance in investigating the ecology of both common and rare tick species. Information on the geographic distribution, host-association, and seasonality of the tick community can help researchers and health-officials to identify high-risk areas.

## Background

The Centers for Disease Control reported a 3.5× increase in vector-borne diseases in the USA between 2004-2016, with 76.5% of cases caused by tick-borne pathogens [1]. The increase in tick-borne disease is attributed to multiple abiotic and biotic factors. Changes in geographic variability and the temporal dynamics of tick species may influence tick-borne disease outcomes. Several endemic tick species have been expanding into new habitats across North America, posing novel risks to local communities [2]. Although there are many tick-borne pathogens, the vast majority of tick-borne disease cases are caused by *Borrelia burgdorferi* [1,3], the main etiological agent of Lyme disease in the USA. Pennsylvania has had the highest number of total Lyme disease cases since 2000, with increasing numbers of annual cases across several counties (Figure 1).

**Figure 1:**
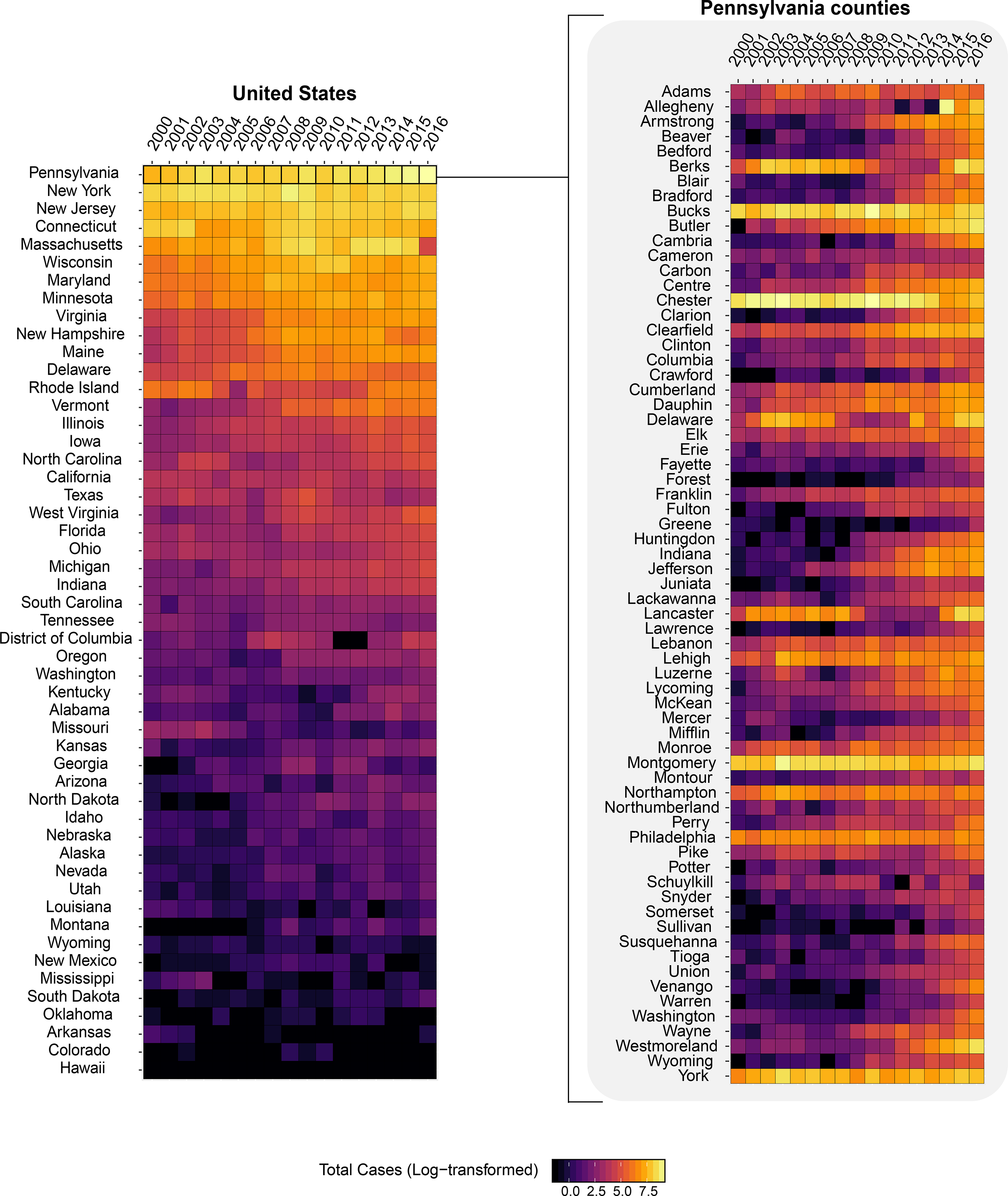
Annual reported cases of Lyme disease by state from 2006-2017 (Left) and the annual reported cases of Lyme diseases by counties in Pennsylvania from 2006-2017. Public data from the Center of Infectious Disease.

Surveillance can be a powerful tool for detection of introduced species (transient or established), emergent arthropod-borne pathogens, and disease risks due to increases or changes in vector community composition. Active tick surveillance approaches such as dragging, flagging, CO_2_-trapping, or live animal capture, can be very effective for assessing tick load by habitat [4,5]. However, active surveillance is labor-intensive, costly, and difficult to implement over a wide geographic area. Passive surveillance, in which citizens submit ticks for identification and/or pathogen testing, is more cost-effective and less labor-intensive and can provide insight into ectoparasite abundance, host associations, or habitat associations across a wider geographic area [6]. Even when tick samples may be in less-than-ideal conditions (e.g. missing taxonomically diagnostic), or data, citizen-submitted tick can accurately represent tick encounter frequency and potential risk of tick-borne disease exposure [7].

Passive surveillance data collected over decades may reveal spatio-temporal changes in ectoparasite communities [8]. Data such as spatial distribution and occurrence of both abundant and rare species of ticks can be correlated with landuse (e.g. habitat loss,fragmentation, management), fluctuating environmental conditions, or changes in human or animal behavior (e.g. encroachment may bring reservoir hosts such as groundhogs in closer proximity)[9,10]. Additionally, long-term surveillance data can also reveal shifts in temporal dynamics of tick populations and communities [2]. The seasonality of known tick species have been described, but year-to-year distribution of tick species may be influenced by inter-annual variability (e.g. local climate) and biotic factors (e.g. local reservoir species abundance). These data can be used to create predictive models that accurately measure risks of tick-borne zoonotic agents.

We present a retrospective analysis of tick collection data in Pennsylvania from the early 1900s to June of 2017. Some of the data prior to 1968 had been published in a USDA report on ticks from Pennsylvania, but were presented in a format that included anecdotes and overall percentages rather than raw number breakdowns by species (Snetsinger 1968). In this manuscript we revisit these specimens and utilize the raw data from both these specimens (1900-1960s) as well Pennsylvania ticks submitted from 1968-June 2017 to identify shifts in tick community composition, phenological patterns, and host associations. We used our database to map the distribution of major tick species at the county level, investigate tick community spatiotemporal dynamics, and explore host and vegetation associations by tick species.

## Methods

### Study locations

The state of Pennsylvania (PA) is located in the mid-Atlantic region of the United States (latitude: 39° to 42°N and longitude: −80° to −74° W). The climate varies across Pennsylvania depending on the region and altitude, but it typically consists of hot, humid summers and cold winters with heavy snowfalls in certain areas. The majority of Pennsylvania’s land-use is dedicated to agriculture (both croplands and pastures), forestland, and dense urban areas. There have been significant changes in the human populations of PA from 1960 to 2010, but a large proportion of the PA population has remained heavily clustered around Philadelphia and Pittsburgh, which are located in south-eastern and in south-western PA respectively (Figure 2). While most specimens were collected within state boundaries, a few were declared from people either visiting or returning from visiting other states. Tick specimens identified as species that are not commonly found in Pennsylvania were later discovered to have been imported or on exotic animals. These non-PA data were not included in state-wide analyses, but were included in the Supplemental Tables (Table S1).

**Figure 2:**
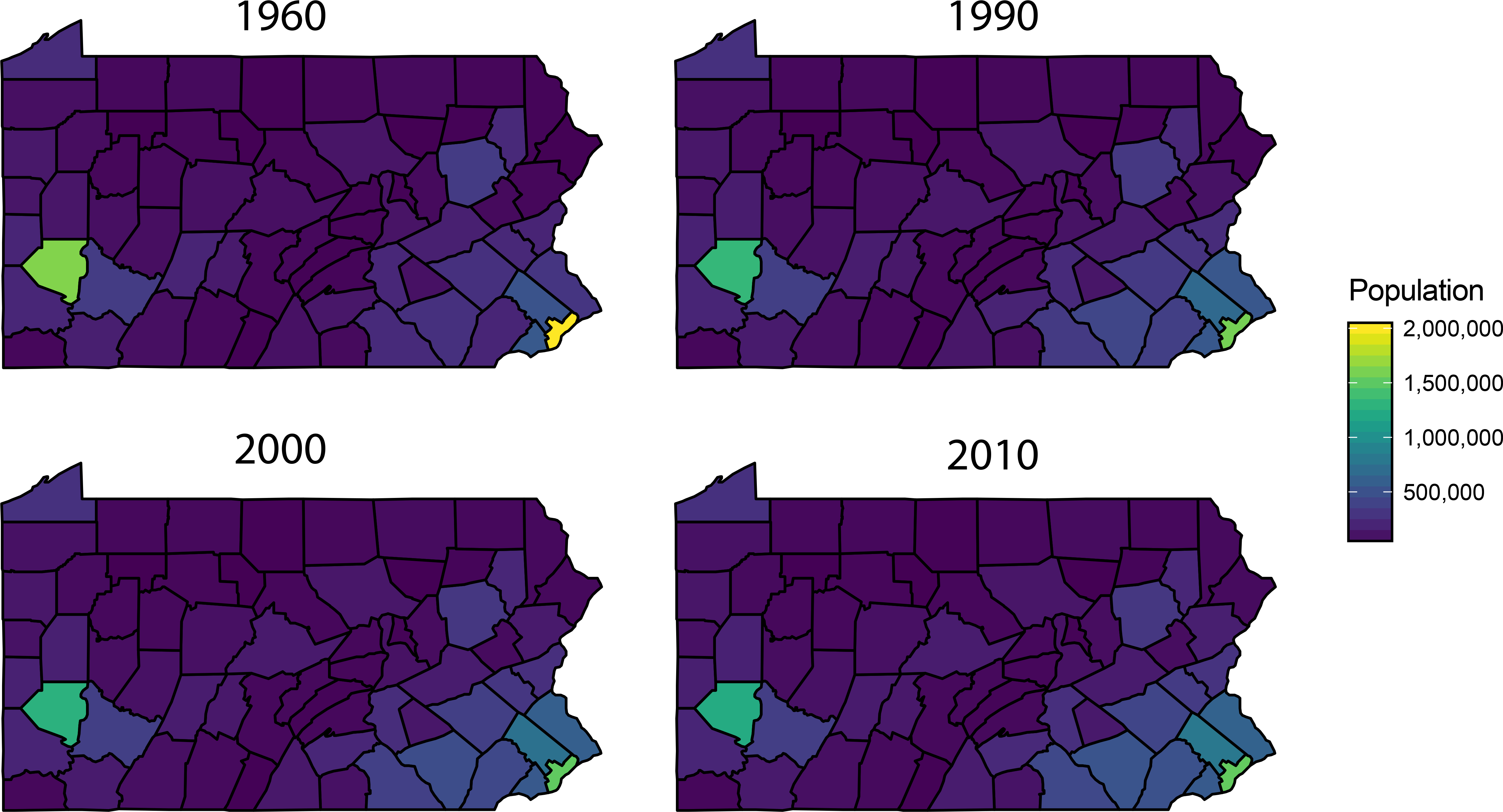
Total population in Pennsylvania counties in 1960, 1990, 2000, and 2010.

### Submissions

The PSU Frost Entomological Museum (hereafter ‘Frost Museum’) houses arthropod samples collected by researchers, teaching collections, and samples submitted by the public for identification. Although some of these samples date as far back as the late 1800s, we only present our analysis of the tick specimens from 1900 to June 2017. Because tick samples were submitted to the Department of Entomology or the Frost Museum over a period of 117 years, they represent multiple collection/submission periods (early 1900-1959, 1960s, 1970-1988, Tick Research Lab (TRL) submissions from 1990-1993, and 1995-present). Two public campaigns account for the majority of the specimens. The first campaign (between 1963-1967) was conducted by Dr. Robert Snetsinger. Dr. Snetsinger collected approximately 500 ticks using a combination of dragging, sweeping, live animal trapping, and roadkill examinations of mammals and birds to assess tick abundance in localized areas (Snetsinger, 1968). He enlisted the help of the public through advertisements in radio, television, and newspapers to obtain 700 additional specimens (Snetsinger, 1968). A second funded campaign dedicated to estimating tick abundance and species diversity was launched by Steven B. Jacobs (second author) from 1990-1993 (TRL). In total 1357 submission lots made up of 3561 tick individuals were contributed by members of the public or by physicians requesting identification.

Tick submitted prior to 1960 reflect the sporadic nature of citizen-based submissions. An active advertisement campaign was used to enlist the help of citizens in the 1960s and again in the early 1990s. Subsequently, post-funding submission rates decreased in volume, but tick submissions are still received regularly by PSU Entomology for identification. Data from the ticks submitted prior to 1989 were identified by museum staff and stored in alcohol at the Frost Museum. Submissions from the general public received from 1989 until June of 2017. TRL submissions were identified by SBJ and stored in alcohol. All data were combined into a single dataset for our analyses. We compiled these data to identify how species abundance have changed over the years. We defined a submission lot (‘submission’) as a vial or lot containing one or more ticks. For analysis of host-association, we used “submissions” versus total tick counts by host. We chose to use this more conservative measure rather than the numbers of ticks submitted to avoid a skew in abundance by host. For example, a submission lot of 1 tick versus 50 ticks from a host were both treated as one ‘submission’. For analysis on the distribution of tick species over time, we used total tick counts.

### Identification

Ticks were morphologically identified to species and life stage using the following taxonomic keys: Argasidae, Ixodidae east of the Mississippi, *Dermacentor*, nymphal *Ixodes*, and nymphs of *Amblyomma* [11–17]. Identification of samples to species is crucial since at least 3 Dermacentor species, 3 species of *Amblyomma*, and 9 different *Ixodes* species have been reported in Pennsylvania. If diagnostic characters were missing due to damage to the specimen, the next level of taxonomic identification was used (*e.g.* samples with missing mouthparts that were clearly Prostriata were identified as “*Ixodes spp*.”). In a few cases, samples were not identified beyond “tick” and were designated “Ixodidae” for hard ticks or “Argasidae” for soft ticks. Difficult-to-identify, or unusual specimens were sent to the National Tick Collection, Georgia Southern University for confirmation (by Dr. James Oliver at the time of confirmation).

### Spatial distribution of ticks

To get an overview of the spatial variability of the tick community from 1900 to 2017, we summed the total tick counts by species across the counties. Rarer tick species with less than 150 individual specimens were grouped by genus. We then plotted the total individuals with a dot-density map with each dot representing a single individual. As the associated geographic data for each submission was at the county-level, the placement of the dots within each county’s boundary was randomized.

We focused on the geographic distribution of the five most abundant species in our database, which are of significant public health and veterinary importance: *Amblyomma americanum (Linnaeus), Dermacentor variabilis (Say), Ixodes cookei (Packard), Ixodes scapularis (Say), and Rhipicephalus sanguineus (Latreille)*. Working on the assumption that counties with higher populations would submit more specimens than less populated counties, we estimated the incidence rate (the total numbers of individual ticks per 100,000 people). This was done by adjusting the total tick count of each species by the county's total population. Because we did not have the county population data for each year, we looked at relevant time periods during the surveillance program: 1960-1970, 1990-2000, 2000-2010, and 2010-2020. For each time periods, we used the 1960, 1990, 2000, and 2010 United Census data respectively to calculate the incidence rate (Figure 2) (U.S. Census data from 1960 to 1990: retrieved from <https://www.census.gov/population/cencounts/pa190090.txt>. 2000 U.S. Census data: U.S. Census Bureau; Table DP-1, Census 2000, Profile of General Demographic Characteristics Summary File 1 using American FactFinder <http://factfinder2.census.gov>, Table retrieved: July 1, 2017). 2010 US Census data: U.S. Census Bureau; Table DP-1, Census 2010, Profile of General Demographic Characteristics Summary File 1 using American FactFinder <http://factfinder2.census.gov>, Table retrieved: July 1, 2017).

For the tick species with less than 150 submissions across 1900 to 2017, we aggregated the submissions by genus. We excluded the five major tick species (*Amblyomma americanum, Dermacentor variabilis, Ixodes cookei, Ixodes scapularis, and Rhipicephalus sanguineus*) from this analysis. We then mapped presence or absence of each genus by county.

### Temporal analysis

To investigate how the annual submissions changed over the course of the passive tick surveillance program, we first aggregated all individual tick specimens by year. Because there were few submissions in the beginning of the program, we grouped the annual submissions into decades starting from 1900-1910 (the years included would be from 1900 to 1909) to 2010-2020.

We analyzed the temporal dynamics of the five most abundant taxa (*A. americanum, D. variabilis, I. cookei, I. scapularis*, and *R. sanguineus*). We did not evaluate total counts by year as it varied drastically during active campaigning for citizen submissions or introduction of identification fees. Therefore, we looked at the proportional contribution of each species to the annual summed counts of the five major species. To detect if there have been any monotonic trends (*i.e*. gradual shifts in abundance), we ran a non-parametric, two-sided Mann-Kendall trend test on the yearly proportion of each of the species from 1900-2017.

### Seasonality

For analyzing seasonal patterns of submissions in the passive-surveillance program, we looked at the months of when the tick specimens were received for identification. For some submissions, there were no dates of when the ticks were found by the citizen, so we used the date that the specimens were received. We first looked at the frequency of the months in which the ticks submissions were received for the collective tick community across 1900-2017 and then by decades. We then investigated the seasonality of the five most abundant tick species individually. For some of the submissions, information of the life-stage (larvae, nymphs, and adults) was included. We then explored if the distribution of the monthly submissions for the life-stages for the major tick species.

### Associated data

Where available, we utilized host-association data for our analyses. Submissions generally included date of tick discovery, location, vegetation associated with tick encounter, from vertebrate the tick was collected (if any), and any additional comments that might provide further insight. The data on human hosts such as gender and age were not included.

#### Vegetation type

For a small subset of these data (1989-1990, TRL) the vegetation associated with the tick encounter were also recorded. We defined the dominant vegetation types as brush, forest (inclusive of mixed, hardwood, and evergreen), pasture (land intended for grazing), managed (residential or urban landscapes), and ecotone (used broadly here to refer to any zone of transition between two plant communities). We used R (Version 3.4.1, RStudio version 1.1.383) to performed a two-way analysis of variance (ANOVA) test to determine whether total tick counts were dependent on the different types of vegetation by tick species.

#### Host Associations

Host information was available for many of the tick specimens (combined by family, except for dog, cat, human, and groundhog). Host data were classified as either domestic or wildlife. We summarized the host-tick data by summing the total submissions by both the tick species and the host groups. We constructed a circular network map to visualize the relationships between tick species and hosts. All host association analyses were done with R (Version 3.4.1, RStudio version 1.1.383) with the packages ‘MannKendall’ for the Mann-Kendall test and the ‘circlize’ package for chord diagrams of host association mapping [18,19].

#### Availability of data and materials section

The dataset(s) supporting the conclusions of this article is(are) available in the passive_tick_surveillance_2018repository, https://github.com/pakdamie/passive_surveillance_tick_2018/tree/master/MAIN_DAT

## Results

### General observations

From 1900 to 2017, PSU Entomology handled a total of 4,491 submission lots consisting of 7,132 tick specimens from twenty-three species (Table 1). There were five species of ticks that accounted for the majority (91%) of the total number of tick: *Dermacentor variabilis* (n = 3172), *Ixodes scapularis* (n = 1899), *Ixodes cookei* (n = 897), *Rhipicephalus sanguineus* (n = 332), and *Amblyomma americanum* (n=196). Other tick species that were represented in at least 100 submissions were *Dermacentor albipictus* (*Packard*) (n = 107), *Ixodes dentatus* (*Marx*) (n = 120), and *Ixodes texanus* (*Banks*) (n= 111). The remaining ticks had < 100 submissions/species and included both hard and soft tick species.

**Table 1:** The total submissions to the PSU Department of Entomology/Frost Entomological Museum from 1900 to 2017. Generic names that have been changed since the submission date are shown in parentheses. Specimens which were not identified to species were listed under the genus and “sp.”

| **Species** | **#Submission lots** | **#Specimens** |
| --- | --- | --- |
| *Amblyomma americanum* | 183 | 196 |
| *Amblyomma cajennense* | 1 | 1 |
| *Amblyomma dissimile* | 8 | 8 |
| *Amblyomma longirostre* | 1 | 1 |
| *Amblyomma maculatum* | 5 | 5 |
| *Amblyomma (Aponomma) transversale* | 2 | 2 |
| *Amblyomma sp.* | 2 | 2 |
| *Argas cooleyi* | 2 | 2 |
| *Argas persicus* | 3 | 3 |
| *Carios (Ornithodoros) kelleyi* | 18 | 18 |
| *Dermacentor albipictus* | 81 | 107 |
| *Dermacentor andersoni* | 1 | 1 |
| *Dermacentor variabilis* | 1498 | 3172 |
| *Haemaphysalis chordeilis* | 1 | 1 |
| *Haemaphysalis leporispalustris* | 38 | 78 |
| *Ixodes affinis* | 1 | 1 |
| *Ixodes angustus* | 39 | 39 |
| *Ixodes cookei* | 661 | 897 |
| *Ixodes dentatus* | 115 | 120 |
| *Ixodes marxi* | 46 | 48 |
| *Ixodes muris* | 6 | 6 |
| *Ixodes scapularis* | 1395 | 1899 |
| *Ixodes texanus* | 16 | 111 |
| *Ixodes sp.* | 79 | 80 |
| *Rhipicephalus sanguineus* | 287 | 332 |
| *Rhipicephalus sp.* | 1 | 1 |
| Not determined | 1 | 1 |
| **Total** | **4491** | **7132** |

#### Spatial Analysis

Ticks were submitted from all 67 counties in Pennsylvania (Figure 3). We hypothesized that more tick submissions would come from areas with higher human populations and as expected, tick submissions were heavily clustered around Allegheny and Philadelphia County where Pittsburgh and Philadelphia are located respectively. When we adjusted the total tick count by county population-decade, we found higher incidence rates in *less* populated counties. For example, in 1990-2000, the highest incidence rates of *Ixodes scapularis* submissions were from Elk County (870 individuals per 100,000 population). Neighboring Forest and Cameron counties also had high submissions of *I. scapularis* with 116.64 and 589.97 individuals per 100,000 respectively. Other counties with high *I. scapularis* incidence rates included Northumberland (299.56 individuals per 100,000), Snyder (198.98 individuals per 100,000), Union (171.31 individuals per 100,000), and Clearfield (99.21 individuals 100,000) (Figure 4).

**Figure 3:**
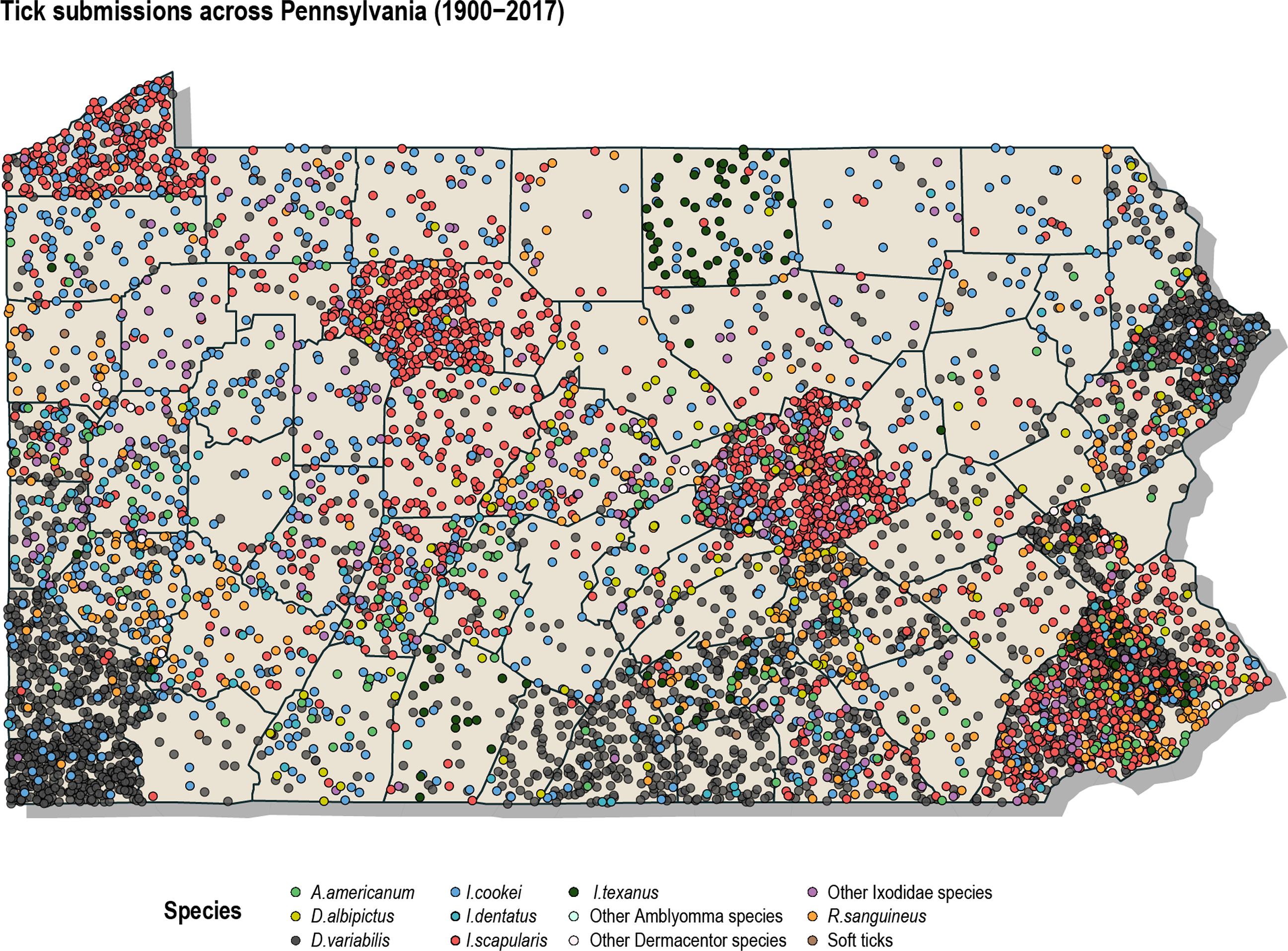
Dot-density map of all individual tick specimens across Pennsylvania from 1900-2017. Each point represents an individual specimen with its placement randomized within the county.

**Figure 4:**
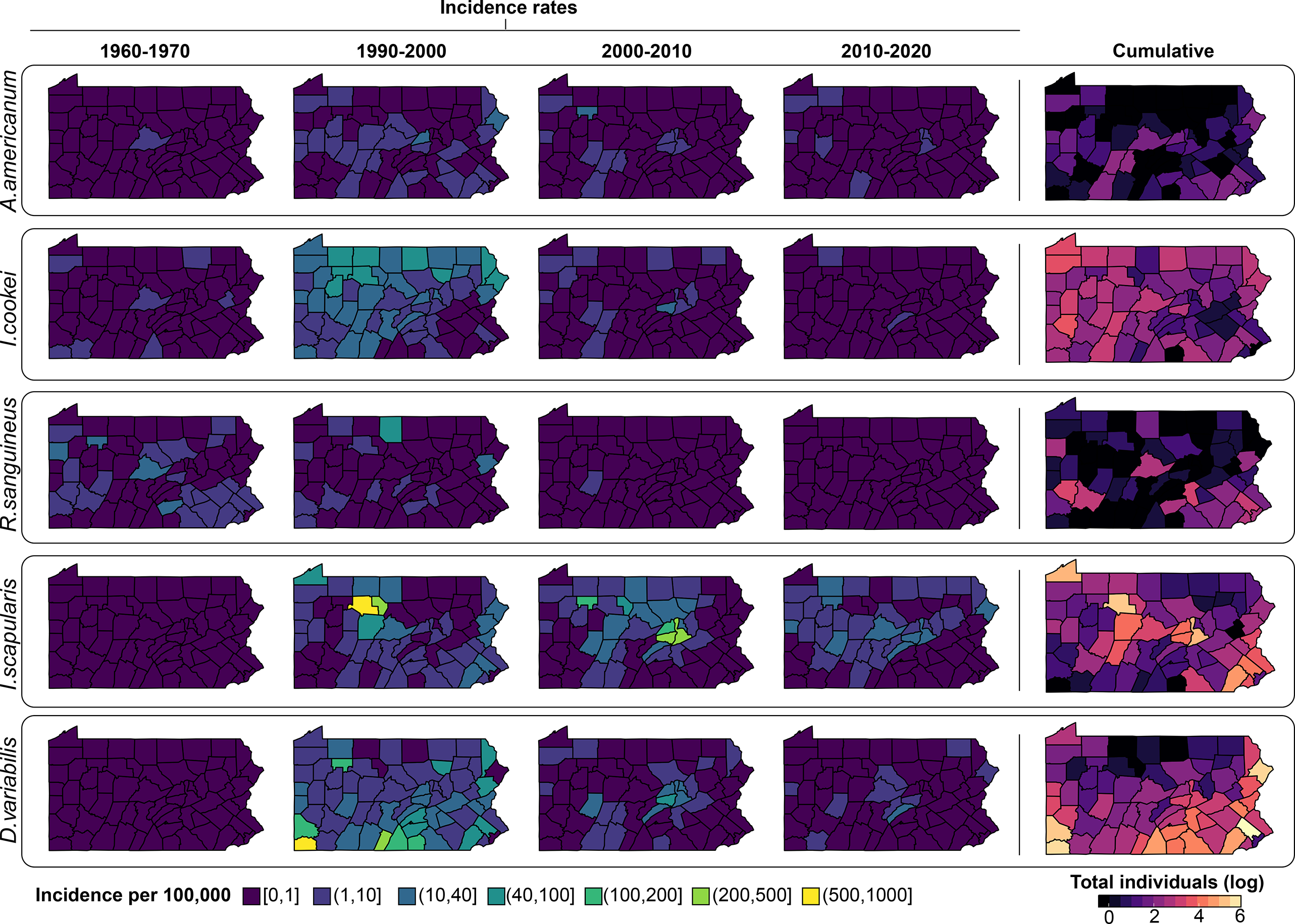
Incidence rates of the five most abundant tick species across Pennsylvania at different time periods from 1960-2018. On the left, is cumulative number of individuals.

*Dermacentor variabilis* distribution was largely localized to southern portions of the state, in 1990-2000, the highest proportion of *D. variabilis* submissions came from Greene County, the most southeastern county of Pennsylvania (865.45 submissions per 100,000). Other southern counties with significantly high rates included Fulton County (350.60 per 100,000) and Franklin County (117.26 per 100,000).

*Ixodes cookei* was more evenly distributed throughout Pennsylvania, although similar to *Ixodes scapularis*, it was more highly abundant in the northern counties. In 1990-2000, Forest County had the highest incidence rates of *Ixodes cookei* with 80.87 per 100,000. *R. sanguineus* and *Am. americanum* had very few submissions and their distribution was mostly scattered across Pennsylvania.

Multiple species within the genera *Ixodes* and *Dermacentor* were widely distributed across Pennsylvania (*I. scapularis, I. cookei, D. andersoni*, and *D. albipictus*) (Figure 5). Other species in the genera *Amblyomma, Argas, Carios (Ornithodoros)*, and *Haemaphysalis* were not as widely distributed, possibly because these species are not commonly encountered or because the specimens were introduced from their native geographic ranges. For example, we only had four submissions of *Argas cooleyi* and *A. persicus*.

**Figure 5.**
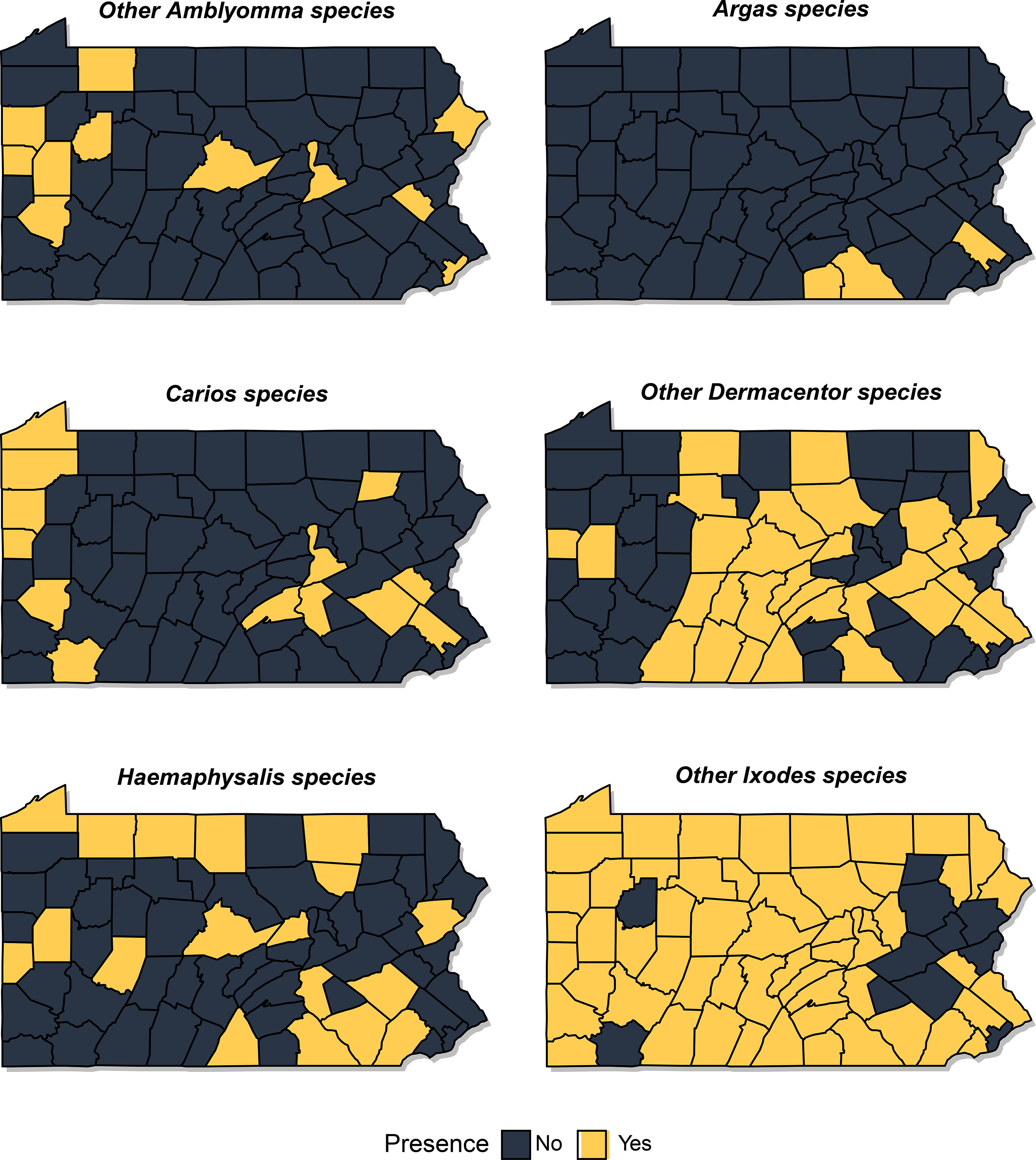
Presence or absence map of the tick genera excluding *Ixodes scapularis, Ixodes cookei, Dermacentor variabilis, Amblyomma americanum*, and *Rhipicephalus sanguineus*.

#### Temporal

##### Temporal shifts in species abundance

Prior to the 1990s, the majority of the tick submissions were identified as *I. cookei* and *R. sanguineus* (Figure 6). The spike in the number of submissions 1990 were largely due to *D. variabilis*, but gradually, *I. scapularis* became the dominant taxon submitted. Results from the Mann-Kendall test supports these observations with an upward trend in the *I. scapularis* counts (tau = 0.288, p = 0.02) and a significant downward trend in *D. variabilis* (tau = −0.408, p = 0.002). The Mann-Kendall also indicate that the proportional contributions of *I. cookei* (tau = −.607, p < 0.001) and *R. sanguineus* (tau = −0.377, p = 0.005) to the total count have also significantly shifted over a century.

**Figure 6:**
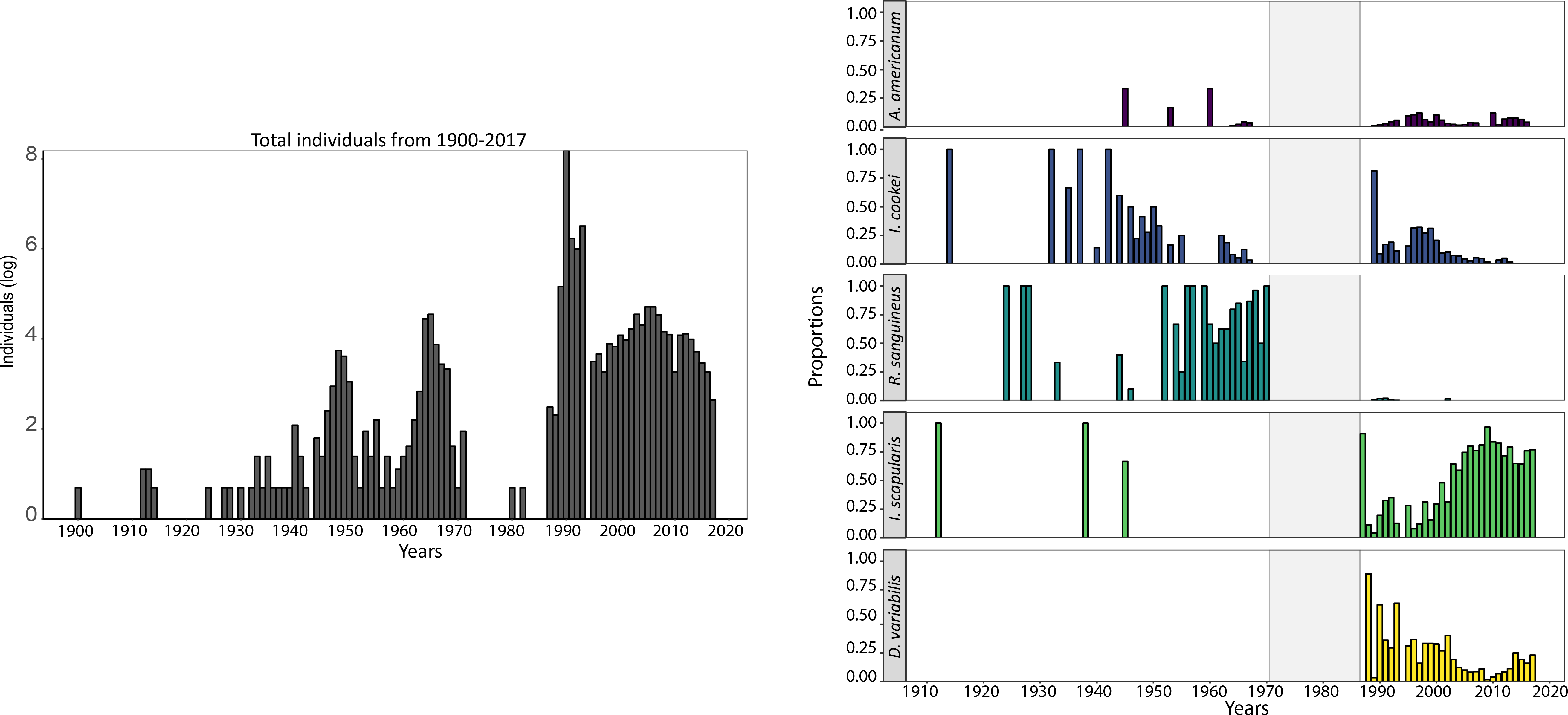
On the left is the annual sum of tick specimens (log-transformed) from 1900 to 2017. On the right are the proportional contribution of the major tick species to the total tick counts (1900-2017). The grey shaded area represent area where there was no tick submissions.

#### Seasonality

Overall, we find that the majority of tick specimens were received in the months between April and July with May being the month with the highest proportion of tick submissions (Figure 7). However, there was significant variation in the seasonality of the submissions during the program (Figure 8).

**Figure 7:**
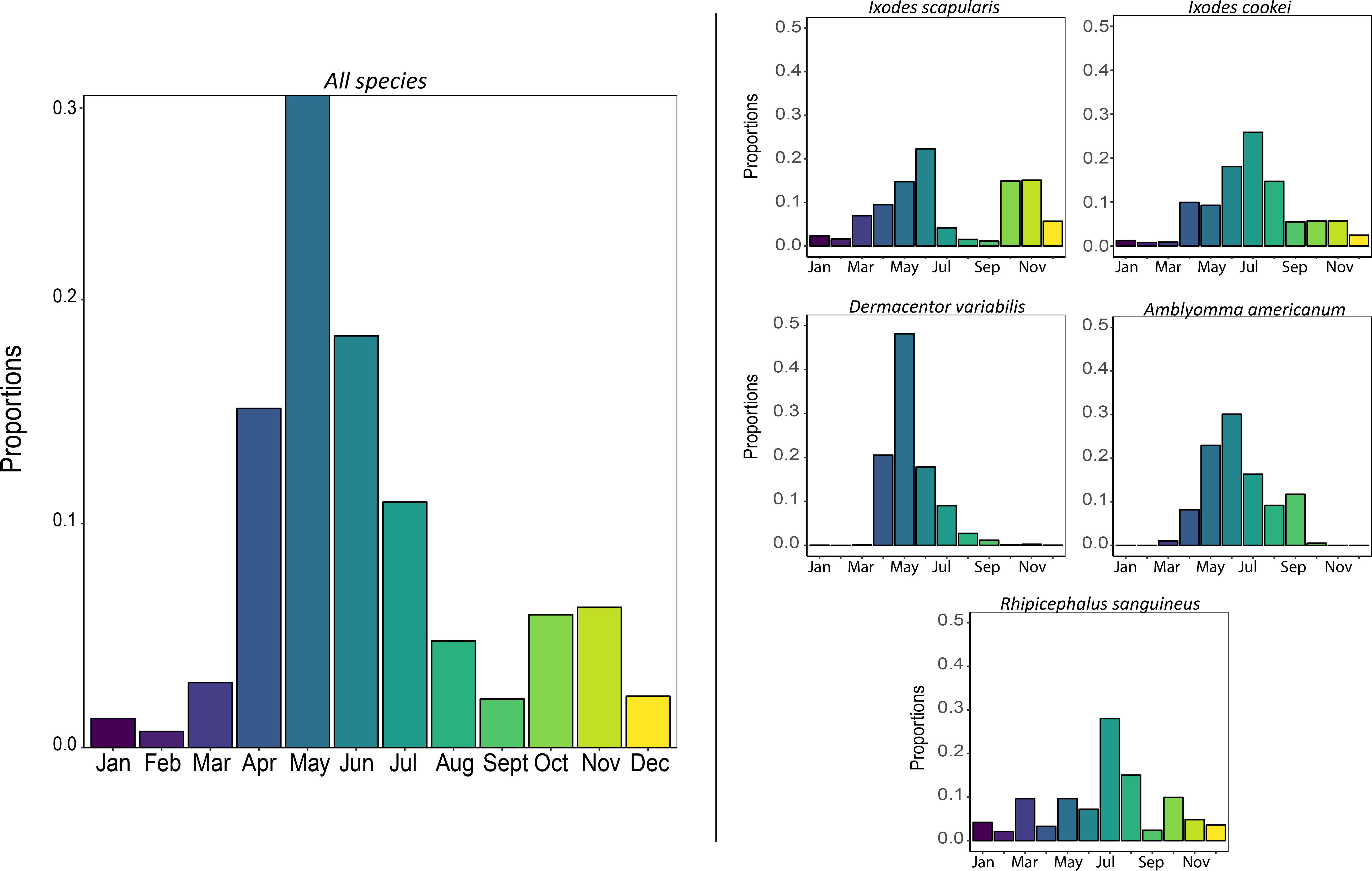
On the left is the total proportion of tick specimens received at different months of the years from 1900 to 2017. On the right are the proportion of the five major tick species received at different months of the years from 1900-2017.

**Figure 8:**
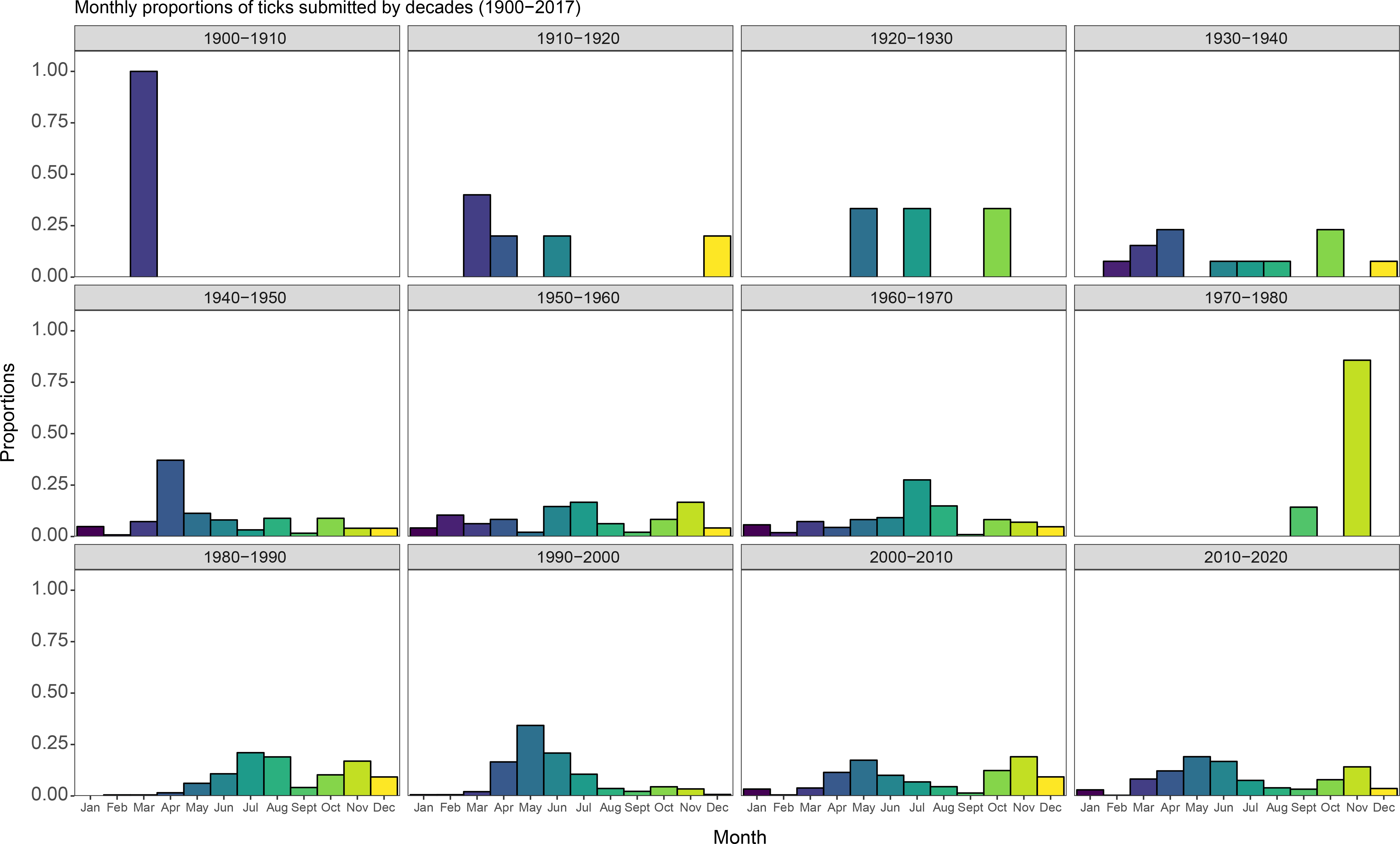
The monthly proportion of the tick specimens received at different months of the years aggregated at different decades.

Submissions of *D. variabilis, Am. americanum, I. cookei* and *R. sanguineus* were most abundant during the period between May and July. By species, *D. variabilis* and *Am. americanum* were most abundant from March to October. *I. cookei* and *R. sanguineus* samples were submitted throughout the year, but had peak abundance in June. Samples of *I. scapularis* were also submitted year-round, but the peak abundances were bimodally distributed, with a large peak in between May to June, and a second peak between October to November. Our data tracks these peaks, although nymphal *I. scapularis* were also submitted during late spring to early fall.

### Life stage abundance by species

There were 6,233 specimens for which the lifestage data were available. Four percent of the submissions were larvae (n=237), 20% of the were nymphs (n = 1271), and 75% of the submissions were adults (n = 4725).

*For Dermacentor variabilis*, the submissions included 32 larvae, 33 nymphs, and 3059 adults from 1960 to 2017(Figure 9). While we only had larval specimens from 1990-2000, we found that the peak submissions were in September (90%). For the nymphal submissions, there was a more unimodal distribution with the peak centered around June. Finally, the adult submissions of both 1990-2000 and 2000-2010 showed similar patterns with the peak in submissions between May and June.

**Figure 9:**
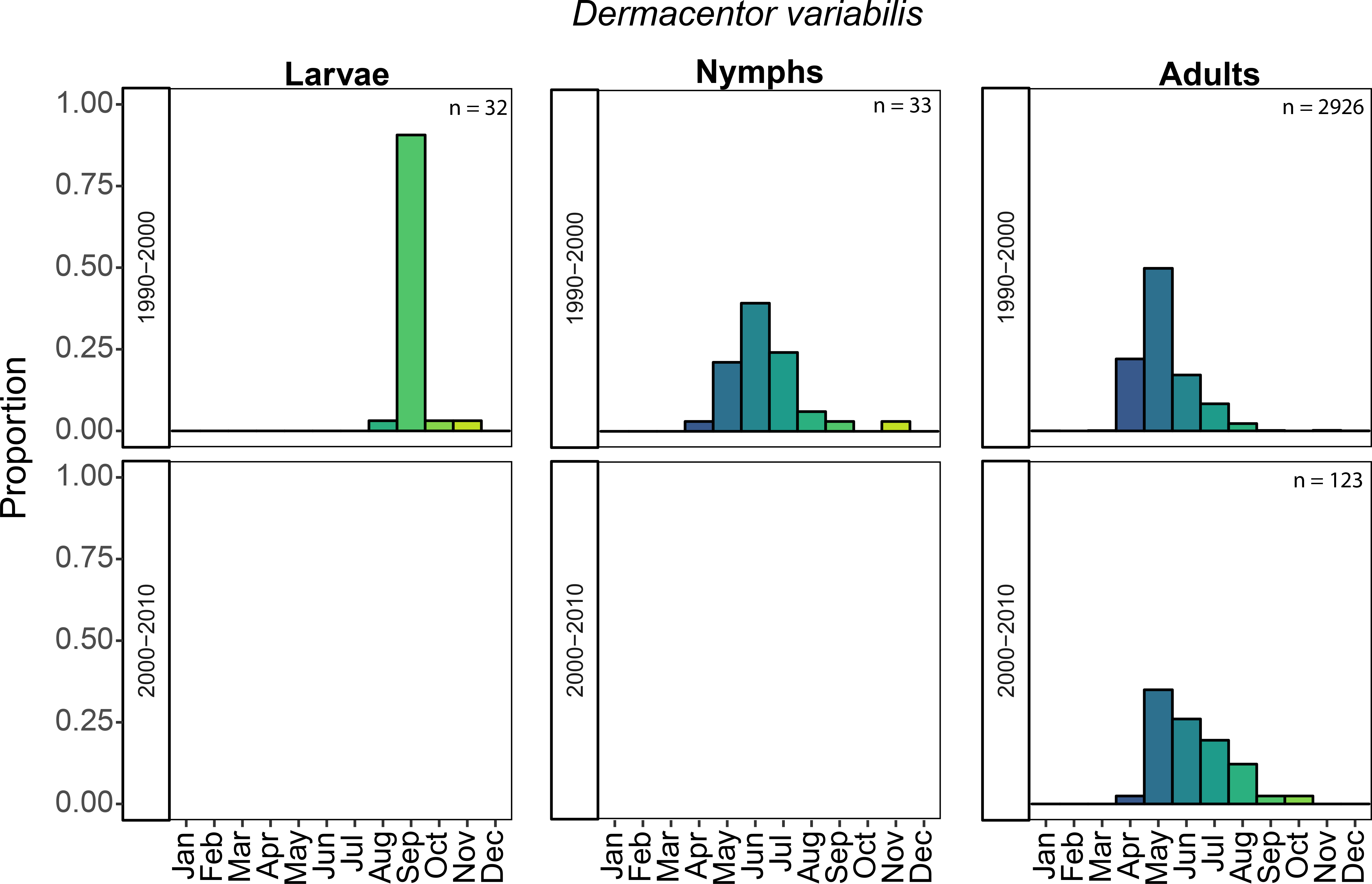
The monthly proportion of *D. variabilis* specimens at the larvae, nymphal, and adult stages across 1960 to 2010

Before 1990, there were only six submissions of adult *Ixodes scapularis* (*Figure 10*). In 1990-2000, the nymphal and larval submissions show a unimodal pattern with the highest proportion of submissions received in June. For the adult submissions during this decade, there are prominent bimodal peaks in May and October with similar proportion of submissions received in both seasons. There were fewer submissions in 2000-2010 with only 23 larvae, 2 nymphs, and 486 adults. The monthly submissions of both the larvae and adults in this decade were consistent to the seasonal patterns found in 1990-2000. Finally, in 2010-2020, the adult submissions (n = 31) show a shift with a number of submissions found earlier in March.

**Figure 10:**
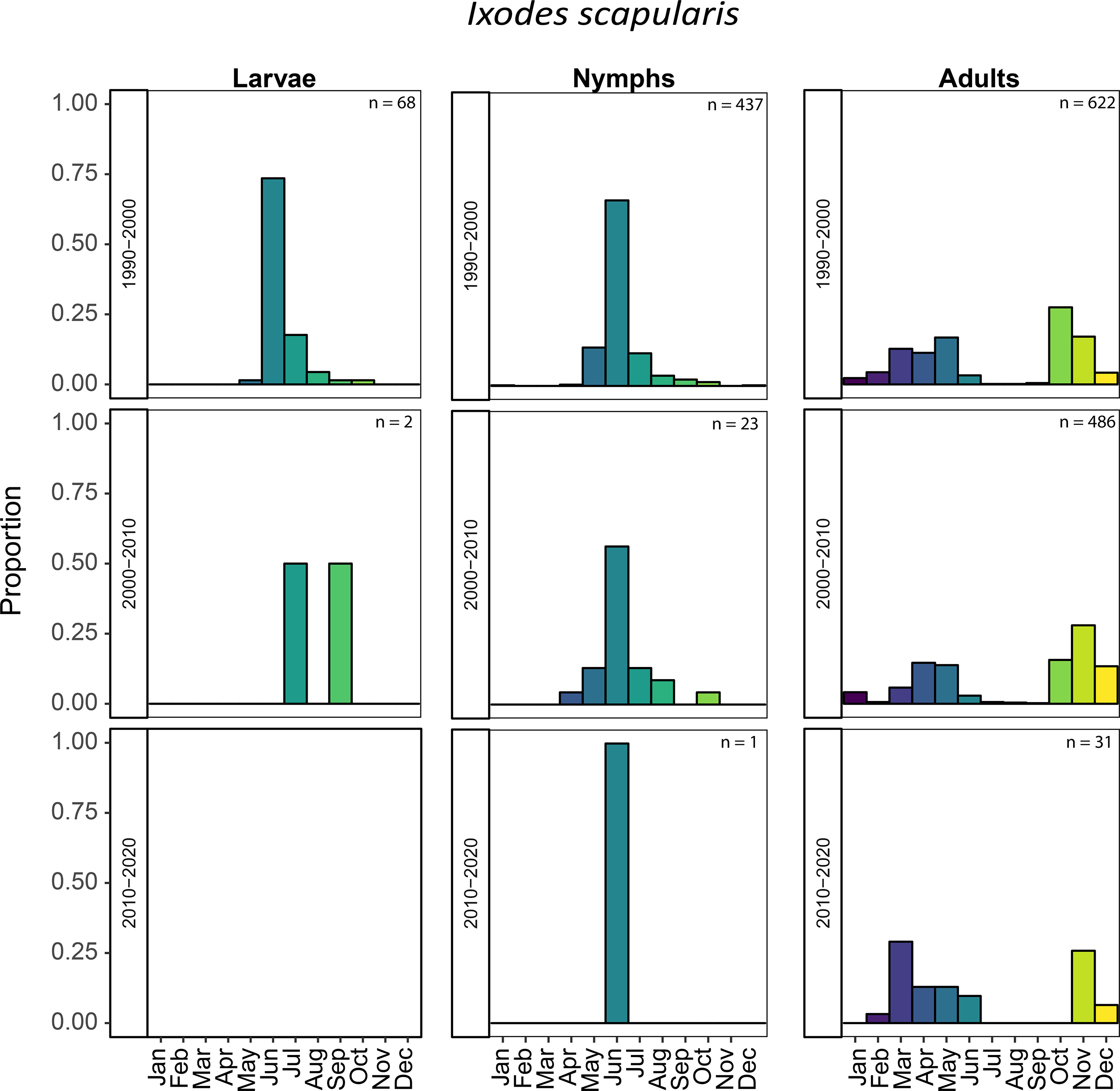
The monthly proportion of *I.scapularis* specimens at the larvae, nymphal, and adult stages across 1990 to 2017

The majority of *Ixodes cookei* submissions were identified as nymphs with a total of 521 submissions followed by adults (n = 182) and larvae (n= 88) (Figure 11). Submission patterns indicate that *Ixodes cookei* specimens are found all year-round specifically in the nymphal stages (Figure 11). Across all life-stages, we see that the distributions are unimodal with peaks in early summer between May and June.

**Figure 11:**
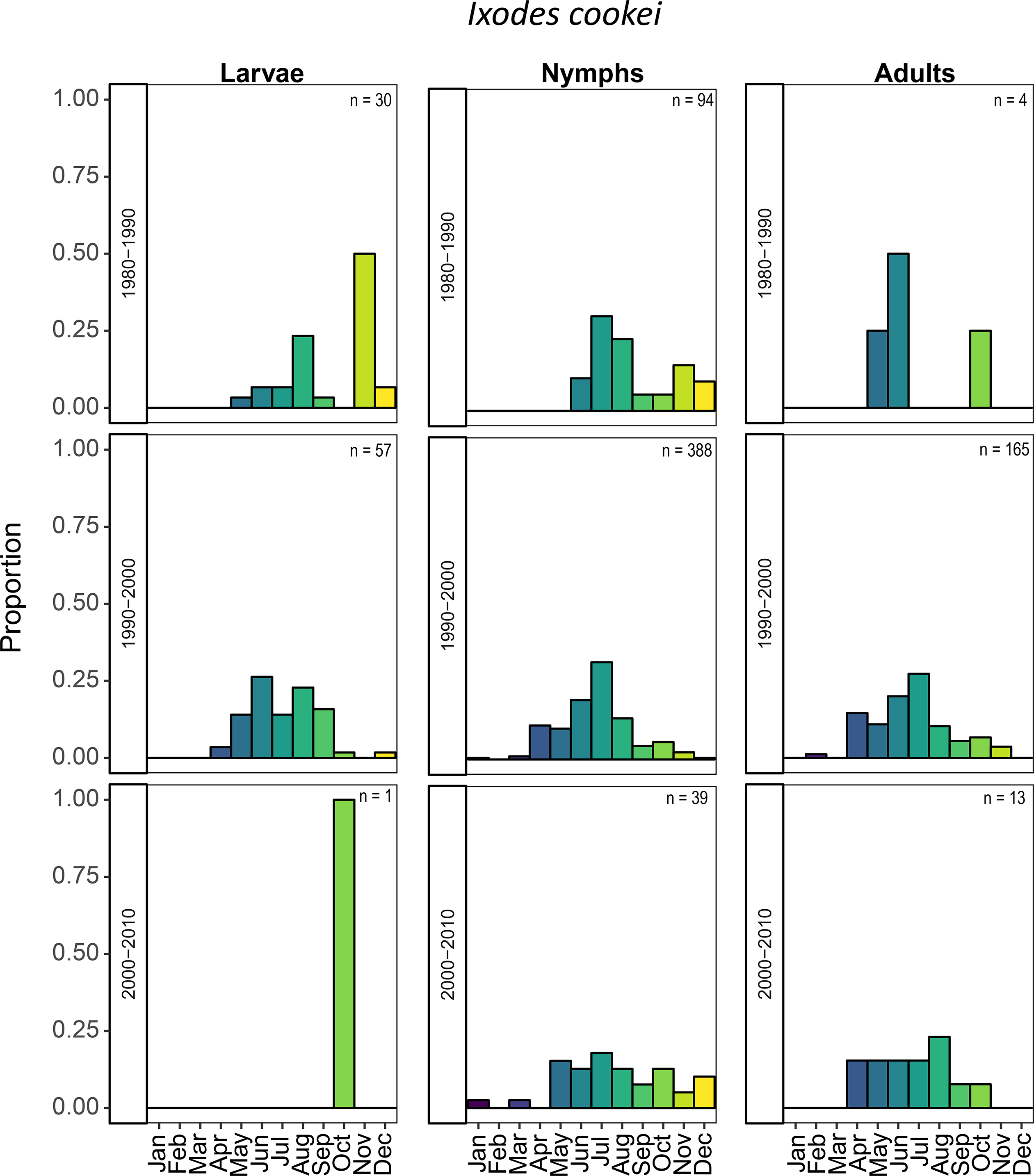
The monthly proportion of *I. cookei* specimens at the larval, nymphal, and adult stages across 1980 to 2017

There were a total of 154 of 183 specimens identified to lifestage for *Amblyomma americanum* (17 larvae, 64 nymphs, and 73 adults). Between 1990 and 2010, the nymphal and adult submission patterns were similar with peak submissions occurring between June and July.

The only decade with high enough counts of *Rhipicephalus sanguineus* to evaluate seasonal variation was the 1960s. In total, we received 59 adult submissions, 17 nymphs, and no larvae.

Submissions were mostly from July and August. The nymphs were submitted in October and adults were submitted between February and September.

#### Vegetation type

There were 677 submissions for which we had data on tick associations with vegetation type (from 1989 to 1990). The large proportion of vegetation data was of ecotone (n = 190) which made up 27 percent of the data. This was followed by forest (n = 178) and managed (n=176). ANOVA f indicated that habitat was a significant predictor for the tick counts (F =3.1997, p = 0.007). We found that ecotone had a weakly significant correlation with tick abundance (F= 2.097, p= 0.036*) and using a pairwise Tukey comparisons in the vegetation types, the greatest difference in the vegetation type was between managed and ecotone.

#### Host association

One of our assumptions about passive surveillance is that there is an inherent bias toward humans as the hosts, particularly since most specimens submitted by the humans on themselves, their pets, or other domestic animals. Figure 12 is a visual representation of the quantitative data on ticks associated with different hosts. By far the majority of submissions were associated with humans and their domestic animals and this reflects the fact that many of the specimens in our collection were submitted by people on themselves or their pets (Figure 12). Of 4491 submissions from PA, there were 2662 attached to humans, 666 were associated with cats or dogs, 20 from other domestic animals, and 168 were submissions pooled from multiple hosts (mixed). There were 11 additional submissions found on various exotic animals. There were 689 submissions for which there was no host record or the ticks were not attached to a host. The remaining 275 were found on various wildlife.

**Figure 12:**
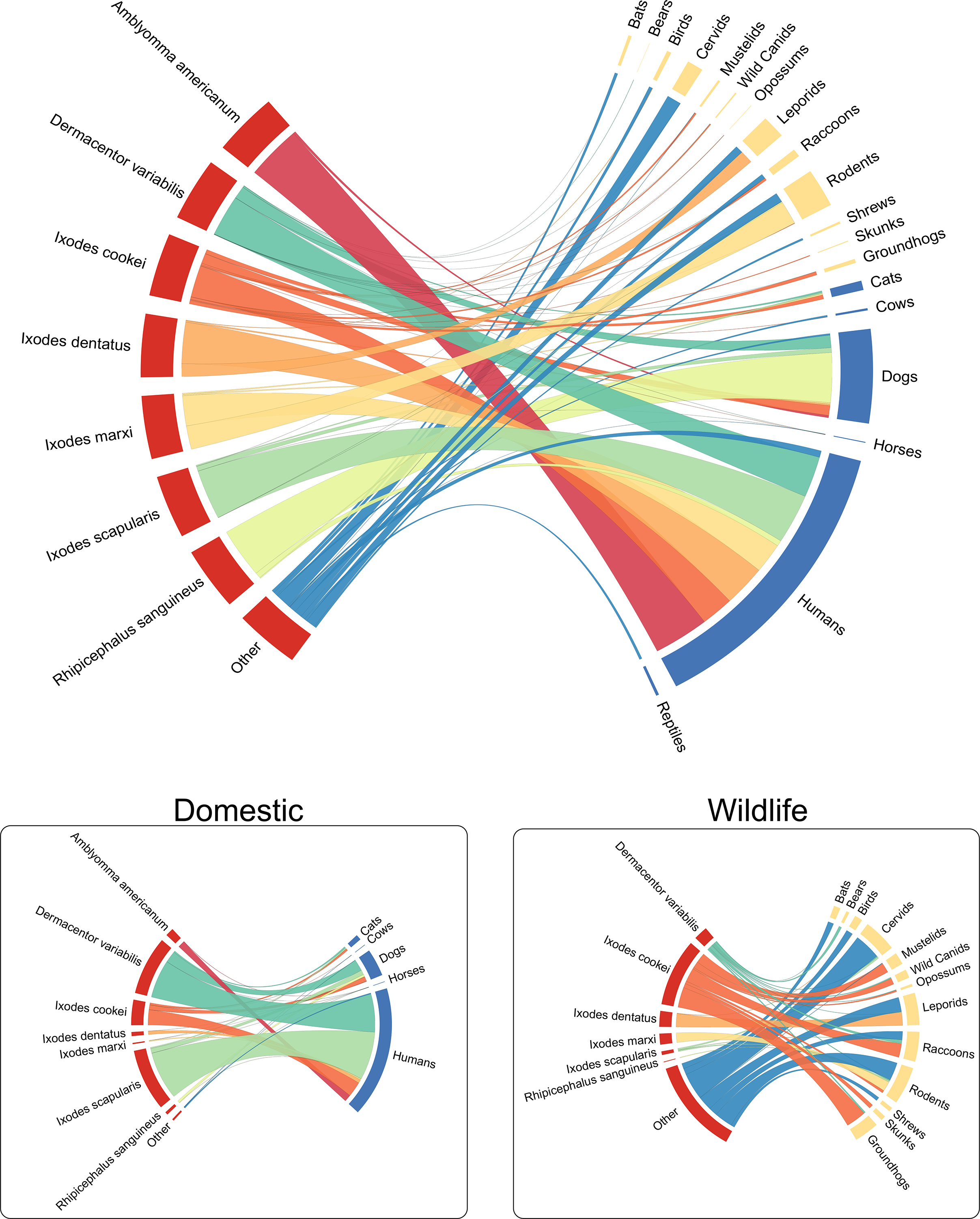
Visual representation of the quantitative data on ticks associated with different hosts.

## Discussion

Our data adds to the current understanding of tick community composition and spatio-temporal dynamics over 117 years. With the caveat that our dataset contains gaps in years of submissions, we were able to detect shifts in tick community composition, seasonality, and host associations that have not been well-documented in a quantitative manner. To our knowledge this is the first time that these data have been compiled in their entirety and analyzed in this format. Since 1993 (~25 years) there have been 28 publications on ticks from Pennsylvania, and 22 of them were focused on *I. scapularis* and/or the microbiota (mostly on pathogens)[20–48]. Subsets of our data had been reported as percentages or combined with data from other museums and literature reviews to assess known distribution of one or more tick species across the state of Pennsylvania [43,49].

Without any data for comparison of other tick species, we can only speculate why there were shifts in the abundance of certain tick species. For instance, *I. cookei* abundance was greatest prior to the 1990s, but has since declined our submission dataset. In contrast, the abundance of *I. cookei* in the Maine passive surveillance program has been constant, even as *I. scapularis* submissions have increased [8]. It is possible that Pennsylvania *I. cookei* abundance has also remained relatively stable, but perhaps we lack sufficient power to detect *I. cookei*.

Apart from one recent fine-scale analyses on the phenology of *I. scapularis* by Simmons et al [42], much of the literature on PA ticks reflects presence/absence data of *I. scapularis* and/or the pathogens they harbor. There have been 2 exceptions: a study of river otters that found 3 female, 1 nymphal and 1 larval *I. cookei* on 3 otters, and a study quantifying the ectoparasites of bats in Blair County, PA that identified 5 specimens of soft tick *Ornithodoros kelleyi* Cooley & Kohls [40,28]. Certainly *I. scapularis* warrants the scrutiny it has received, but other tick species have not been studied closely enough to detect the shifts in tick biodiversity, the potential causes, or epidemiological consequences of these shifts. Further, neither expanding tick ranges nor presence of introduced or established species have been closely monitored. Thus, although we have only recently discovered the presence of the newly invasive longhorn tick (*H. longicornis*) in Pennsylvania, we do not know whether it had been introduced previously. *H. longicornis* is a tick with a wide host range, potentially capable of harboring and transmitting multiple pathogens, may induced meat allergies, and may be capable of reproducing parthenogenically [50–52].

### Temporal comparisons of tick community compositions

In the 1960s, PA tick communities consisted predominantly of three species: *I. cookei, D. variabilis*, and *R. sanguineus*. The most abundant species at that time, *I. cookei*, is often referred to as a groundhog tick, but has been known to multiple mammalian hosts and will readily bite humans and dogs [53]. The second most abundant species, *D. variabilis*, was widely distributed and eventually became the dominant species submitted over *I. cookei* in the 1990s. By 1991 *D*. variabilis had been identified from all but 4 counties [43]. After 1995, *D. variabilis* annual submission rates declined as *I. scapularis* submission rates increased. Although we cannot directly infer a causal negative relationship between these two species from our data, this pattern was also observed in neighboring Ohio. The Ohio State passive surveillance program (started in 1978) did not detect *Ixodes scapularis* (=formerly *I. dammini*) until 1989 [54]. At that time, the dominant species were *D. variabilis* (~97% of submissions) and *Ixodes cookei* (1.2%)[54]. Between 1989 until 2008, *I. scapularis* accounted for less than 1% of the total submissions, but after 2009, the abundance began to increase until in 2012, they accounted for 24.8% of ticks submitted to the Ohio Department of Health [55].

The fourth most abundant species of note in 1968 was *R. sanguineus*, the Brown Dog tick. *R. sanguineus* originated in Africa, but has since become a cosmopolitan urban pest species found worldwide in association with humans and their canine companions [56]. Snetsinger suggested in 1968 that *R. sanguineus* had established breeding populations in Pennsylvania, but according to our records, we have had very few submissions since then and none since 2002. Since both *R. sanguineus* and *D. variabilis* are competent vectors of Rocky Mountain Spotted Fever (RMSF), the recognition of either of these species is important for diagnosis and treatment after a bite (ProMed-mail. Rocky mountain spotted fever - USA (03): (PA) fatal. ProMED-mail 2014; 8 Sept; 20140908.2757657.http://www.promedmail.org July 2018 [57].

The last species that accounted for more than 150 submission lots was *Am. americanum*. Although this species was not common in the 1960s, there was an increase in submissions from 1990s-2000s. Springer et al used predictive modeling to determine that the southern border of Pennsylvania encompassing the Appalachia Mountains were unsuitable for supporting *Am. americanum*, but that western Pennsylvania might experience expansion from the west [44]. We have identified specimens from as recent as October 2016 in our dataset, although we cannot say for certain whether this specimen represents transient introductions or established colonies since the lifestage was not available. *Am. americanum* is a competent vector of ehrlichiosis, tularemia, RMSF, multiple *Borrelia* species (Including *B. lonestari, B. andersonii*, and *B. americanum*), and competent vector of heartland virus)[2,58,59]. *Am. americanum* has also been implicated in inducing meat allergies in certain people [60].

One additional species of note is *Haemaphysalis leporispalustris* (*Packard*) (=*Hlp*), a congeneric of *H. longicornis*. This species abundantly distributed throughout North America [49,53]. *Hlp* is often commonly referred to as a rabbit tick, although it will readily parasitize any mammal or bird and has been known to parasitize humans (Parker et al., 1951; Snetsinger, 1968; Freitas et al., 2009). It can be of public health concern because of its potential role as a vector of Tularemia and Rocky Mountain Spotted Fever [49,62]. In 1968 this species was widely distributed throughout the Pennsylvania, but we have no records of this species after 1993. Since the submissions for tick identification after 1990 were mostly limited to ticks removed from humans or their domestic animals, there is insufficient power to detect this species. Farther north and west, *Hlp* is still found parasitizing a wide range of avian and mammalian hosts [63,64].

The seasonality data for the five most abundant tick species inferred by our passive surveillance data is consistent with previous records of seasonality described by other researcher [5,42,65,66], demonstrating that these types of passive data contain biologically meaningful signal.

### Quantifying host association

Vector-host associations are important factors for predicting risk of pathogen transmission and identifying key players in a sylvatic disease cycle. Common names such as the rabbit tick, deer tick, or dog tick are often used when describing ticks to the public, but can sometimes lead to misinformation about host preferences. For example, many tick resources for the public refer to *I. cookei* as the “Groundhog” or “Woodchuck tick” (e.g. CDC https://www.cdc.gov/ticks/tickbornediseases/tickID.html). Yet, previous literature on *I. cookei* clearly describes it as a cosmopolitan species that parasitizes a wide range of medium-sized mammals [53,67]. Our data support the observations that this species can be found parasitizing human, domestic animals, and wild animals. Although *I. cookei* is important as a vector is of the very rare, but potentially fatal Powassan Encephalitis Virus, it is not a competent vector of *B. burgdorferi* [68,67,69]. It therefore remains a sometimes ignored species for which little is known about its microbiota (including pathogens, commensals, or symbionts). Because of its broad host range and co-parasitism with other tick species, *I. cookei* may be a potential bridge vector that can transmit pathogens from one reservoir host to another.

Host associations of more rarely encountered tick species can sometimes lead to incorrect assumptions about host preferences. While some tick species may be presumed to hold strict host preferences, they may bite humans if given the opportunity. For instance, *I. dentatus* biting humans in cabins that had been inhabited by their squirrel hosts in Maine and Vermont, and *I. marxi* was found (Hall et al., 1991, Lubelczyk et al 2010). In our dataset so-called “generalist” tick species (*D. variabilis, I. scapularis, I. cookei*) were found parasitizing a wide range of vertebrate hosts. Similarly, we found that ‘specialist’ tick species (*I. dentatus, I. marxi,I. muris*) that were mostly associated with a single host or limited to host size (*e.g*. small mammals or birds). Some species known to be multi-host species, however, were limited to one host or few in our collections (*e.g. I. texanus* was only found on raccoons, yet it has been found parasitizing small rodents and medium sized mammals [5,70,71].

### Importance of tick species identification

Merten and Durden (2000) described 84 tick species naturally occurring in the United States, 40 species of which will are known to bite humans (11 species of soft ticks and 29 hard ticks)[72]. Nine of the 40 human biting species are classified by the CDC as important vectors of zoonotic disease website (https://www.cdc.gov/ticks/tickbornediseases/tickID.html): 1) American dog tick (*D. variabilis*), 2) Blacklegged tick (*I. scapularis*), 3) Brown dog tick (*R. sanguineus*), 4) *I. cookei* (groundhog tick), 5) Gulf Coast tick (*A. maculatum*), 6) Lone star tick (*A. americanum*), 7) Rocky Mountain wood tick (*D. andersoni Stiles*), 8) Soft ticks (*specifically Ornithodoros spp*.), and 9) Western blacklegged tick (*I. pacificus* Cooleyi and Kohls). In the last 20 years, *Ixodes scapularis* has become the most abundant tick species in Pennsylvania. Distinguishing *Ixodes* from other genera of ticks is fairly simple, but <u>species</u> level identification requires more detailed morphological examination, since there are 6 endemic species of *Ixodes* and 3 exotic species that could potentially be misidentified as *I. scapularis*. More generally, although many tick species are incompetent vectors of *B. burgdorferi*, they may be vectors and/or reservoirs of other pathogens/parasites, or acquire pathogens during co-feeding [5,21,73,74]. It is therefore important to correctly identify tick species, not only for determination of disease risk, but also because the treatments for the pathogens they transmit may differ significantly.

### Finding context for exotic specimens: the hidden gems in retrospective analyses

Our data encompasses a large timeframe and includes several collection periods. While such a rich dataset held much promise, the reality was that cleaning up and standardizing the data into an analyzable format was a laborious task. Additionally, without context, it was difficult to understand how exotic specimens arrived onto Pennsylvania soil. Much of the literature on the distribution of ticks other than *I. scapularis* in Pennsylvania was found in lists published either in before or during 1940s or after the 1990s. Fortunately, we were able to locate a USDA report on ticks and tick-borne disease by Dr. Robert Snetsinger [49]. In it were explanations for the presence of exotic specimens, but also contained were snapshots into the PA tick community composition as well as the public health concerns of tick experts at that time. Although the report did not have actual counts (only percentages) with associated metadata (and in fact contained data collected from other museums as well as references to data from previous papers or unpublished data), we were able to cross-reference the existing tick specimens from this time period with associated data contained in handwritten notes found at the Frost. The Snetsinger report provided a snapshot the epidemiological focus of the time with respect to tick vectors and the pathogens they could spread. For instance, it was known, but not published at that time, that the rare and deadly Powassan Encephalitis Virus (PEV) was present in groundhogs in southern PA (Snetsinger, 1968). *I. scapularis* is now responsible for more arthropod-borne disease outbreaks in the continental US, but in the 1960s *I. scapularis* populations (which had been decimated by deforestation in the early 1900s) had not yet re-established a foothold in most of the commonwealth[49,76]. The tick-borne diseases of concern from 1963 to 1993 were Tularemia (caused by *Francisella tularensis*) and Rocky Mountain Spotted Fever (caused by *Rickettsia rickettsii*) (Snetsinger, 1968; Snetsinger et al., 1993).

In addition to 19 specimens that were from out-of-state, we also identified several exotic tick specimens within Pennsylvania. One specimen of *Amblyomma cajennense* (the Cayenne tick generally limited to neotropic regions) was collected from a capybara in 1913 from the Philadelphia Zoological Garden [49]. A century later, Brazilian researchers documented spotted fever-infected *A. cajennense* on capybaras [77]. Perhaps if this specimen has been maintained well enough, we might be able to extract DNA and assay for spotted fever rickettsial species.

Another specimen, a single European species of tick, *I. ricinus* (the castor bean tick), and sister taxon to *I. scapularis*, was found on lizards in the Pittsburgh Zoo [49]. A third oddity was a reptile-associated tick, *A. dissimile*, known to be imported through the pet trade (snakes) and on research animals. In one humorous, but almost hidden anecdote, one of the tick specimens was described to come from a snake that was used as part of the costume by a “night club ballerina” [49].

Some exotic specimens suggested periodic introduction, but subsequent lack of establishment. In 1967 there was one specimen of the Gulf Coast Tick (*A. maculatum (Koch))* identified after being removed from the ear of a child who had never left Pennsylvania (Snetsinger 1968). At that time, *A. maculatum* was only known from the neotropics and nearctic regions along the coasts of southern Atlantic states, so the presence of this specimen was “a mystery” (Snetsinger 1968). We identified 4 additional submissions of this species in our database since 1968 from 1969, 1990, 1994, and 2004. In the last several decades, the range of this species has expanded westward into the Mid-west and northward into the Mid-Atlantic on migrating birds, including into neighboring states (Sonenshine 2018). Their establishment northward has been attributed to climate change, but because they require high humidity and higher temperatures, what has driven their movement inland and westward is not yet clear. It is possible that successful breeding populations are more tolerant of cooler and/or drier conditions, or alternatively, because they are localizing to warmer and/or more humid areas along riparian ecotones [3]. *Am. maculatum* is a vector of *Rickettsia parkeri*, a mild fever-causing sickness in the Mid-Atlantic states, but its epidemiological significance is amplified should it share its rickettsial load to *A. americanum* through co-feeding on the same host [3].

There were two soft ticks that were most either introduced or very rare. The first is *Argas persicus*, an Old World species associated with poultry [78]. This is a species rarely collected in the United States and it is presumed to have been imported on poultry. The only reported sightings confirmed by Glen Kohls came from California, Georgia, Maryland, and Pennsylvania [78–80]. Our two confirmed instances of *A. persicus* from Pennsylvania chickens came from York and Adams county [78].

The other argasid species found was *Argas cooleyi. A. cooleyi* is mostly associated with cliff swallows in southwestern states of the USA, so humans do not normally encounter them unless they disturb the nests or if the nests are in close proximity to human dwellings [81]. It is therefore unlikely that this specimen is commonly encountered in Pennsylvania unless imported by humans or through bird movement.

### Impact of free versus per-submission charges for tick identification on submission rates

Rates of tick submissions from the public varied depending on the extent of advertisement and whether there was a cost associated with tick identification. In the case of the 1960s Snetsinger campaign, submission rates were high largely due to a multimedia advertisement campaign that targeted principally housewives, although other citizens also submitted specimens during that period. Similarly, during the first year of the TRL campaign, submissions rates were initial high, but the submission rates declined over the next two years. Subsequent lack of funding to support a free tick species identification meant that to identifications had to be done at cost to continue to provide the service. This meant that submissions were mostly limited to those willing to pay 25 dollars per specimen or to medical professionals who removed the ticks from patients. In addition to the cost of tick identification, there was no available funding for either standardized pathogen screening or database management, which greatly hampered efforts to provide a more robust dataset. This is in contrast to states such as Maine, where the state Department of Health was directly involved in collection of tick data and screening for tick-borne pathogens [8]. In just under 2 decades, the Maine DOH program (after engaging in a sustained outreach campaign to encourage public submissions) obtained 24,519 ticks were submitted for identification free of charge. We suspect that the submission numbers for a statewide tick identification program would likewise be high enough to detect lower abundance ticks and potentially detect invading species.

### Multi-faceted approach to tick surveillance

Passive surveillance is sometimes criticized for under-representation of certain taxa or bias toward certain host associations. However, citizen-submitted tick collections can provide valuable baseline data on prevalence and likelihood of tick encounters ([82–84,7]. In fact, there is evidence that passive tick surveillance data is more strongly correlated with reported human cases of tick-borne diseases than active surveillance [7]. A community engagement program that actively recruits ticks submitted by citizens should be coupled with support for a rigorously curated database of tick submission.

While passive surveillance is able to capture a wide geographic range, active surveillance is able to detect fine-scale population estimates and host associations. Utilizing complementary strategies can help fill in knowledge gaps about tick prevalence. For example, in an exclusively passive surveillance-based study, only 17 of 77 Oklahoma counties were identified as having established lonestar tick populations (*A. americanum*) [85]. In contrast, in a study using a combination of retrospective literature review, data compilation of specimens from archival collections, *and* active collection (dry ice, dragging, and flagging) in counties presumed to be free of *A. americanum*, 68 of 77 counties of Oklahoma were identified as colonized [83]. One of the limitations of active collection methods is that while overall population sizes may be high, sampling may not reflect population abundance, particularly if the distributions are spotty. Tick populations are not static and may be highly mobile, depending on the host upon which they alight. Nevertheless, active surveillance can fill in the gaps in tick distribution throughout each state.

Thus, an ideal surveillance program for vector-borne disease epidemiology would utilize both active and passive collection strategies. Both passive and active collections provide complementary data for accurate assessment of tick-borne disease risk that can be combined with reported human cases of tick-borne disease. The metadata associated with both active and passive tick surveillance (assuming it has been curated and well-managed) can provide insight into tick-host association, vegetation, seasonality, and shifts in population structure that can be used for modeling disease risk. Archival tick samples (or their DNA) can be useful for retrospective mining for research on the population genetics of ticks to detect gene flow, host shifts, or on their microbial inhabitants.

## Conclusion

When we combine citizen-based passive surveillance data with museum collection data, we find that hybrid datasets are a powerful tool for mining past ecological and epidemiological events. Many states maintain county records on passive tick submissions to veterinary or medical health officials, but there may be other cryptic collections (and associated data) housed in museums, universities, government institutions, or with private individuals. In the near future, we are contributing our data to those in the literature and with currently unexplored collections into a massive meta-analysis. These cryptic ectoparasite collections in total will provide the basis for exploring hypotheses such as: 1) are shifts in tick populations correlated with increasing human encroachment on natural habitats, 2) what are some phenological reasons for the increase in *Ixodes scapularis* abundance; or 3) if displacement of a dominant tick community species occurs, what are the implications for tick-borne disease risk? We anticipate that participating in such a study will fill in the gaps of knowledge about less well-studied tick species.

## DECLARATIONS

Steve Jacobs is a retired Senior Extension Specialist with the Department of Entomology at Penn State.

## Acknowledgements

The authors would like to thank Dr. István Mikó for critical manuscript review, the public and researchers who have contributed ticks to PSU, The Frost Entomological Museum for access to the collection, and critical suggestions from anonymous reviewers. This work was supported by funds from NSF GRFP DGE1255832 (to support Damie Pak), the Huck Institutes of Life Sciences, and the Penn State College of Agriculture.

## Author Contributions

SBJ compiled that data into a digital database, collected metadata, identified tick specimens from 1989 until 2017, and manuscript revisions. DP generated data visualizations, conducted statistical analysis, and contributed to manuscript writing and revisions. JMS curated the database, directed analyses, and contributed literature review, manuscript writing, and revisions

---

**Figure S1:** The monthly proportion of *A. americanum* specimens at the larvae, nymphal, and adult stages across 1990 to 2017

**Figure S2:** The monthly proportion of *R. sanguineus* specimens at the larvae, nymphal, and adult stages across 1990 to 2017

**Supplementary Table S1:** Specimens submitted from outside of the state of Pennsylvania.

## References

1. Rosenberg R, Lindsey NP, Fischer M, Gregory CJ, Hinckley AF, Mead PS, et al. Vital SignsJ: Trends in Reported Vectorborne Disease Cases - United States and Territories, 2004-2016. MMWR Morb Mortal Wkly Rep. 2018;67:496–501.

2. Sonenshine DE. Range Expansion of Tick Disease Vectors in North America: Implications for Spread of Tick-Borne Disease. Int J Environ Res Public Health. 2018;15.

3. Eisen RJ, Kugeler KJ, Eisen L, Beard CB, Paddock CD. Tick-Borne Zoonoses in the United States: Persistent and Emerging Threats to Human Health. ILAR J. 2017;58:319–35.

4. Bouchard C, Leighton PA, Beauchamp G, Nguon S, Trudel L, Milord F, et al. Harvested white-tailed deer as sentinel hosts for early establishing Ixodes scapularis populations and risk from vector-borne zoonoses in southeastern Canada. J Med Entomol. 2013;50:384–93.

5. Kollars TM, Oliver JH. Host Associations and Seasonal Occurrence of Haemaphysalis leporispalustris, Ixodes brunneus, I. cookei, I. dentatus, and I. texanus (Acari: Ixodidae) in Southeastern Missouri. J Med Entomol. 2003;40:103–7.

6. Oliver JD, Bennett SW, Beati L, Bartholomay LC. Range Expansion and Increasing Borrelia burgdorferi Infection of the Tick Ixodes scapularis (Acari: Ixodidae) in Iowa, 1990-2013. J Med Entomol. 2017;54:1727–34.

7. Ripoche M, Gasmi S, Adam-Poupart A, Koffi JK, Lindsay LR, Ludwig A, et al. Passive Tick Surveillance Provides an Accurate Early Signal of Emerging Lyme Disease Risk and Human Cases in Southern Canada. J Med Entomol. 2018;

8. Rand PW, Lacombe EH, Dearborn R, Cahill B, Elias S, Lubelczyk CB, et al. Passive surveillance in Maine, an area emergent for tick-borne diseases. J Med Entomol. 2007;44:1118–29.

9. Brownstein JS, Skelly DK, Holford TR, Fish D. Forest fragmentation predicts local scale heterogeneity of Lyme disease risk. Oecologia. 2005;146:469–75.

10. Simon JA, Marrotte RR, Desrosiers N, Fiset J, Gaitan J, Gonzalez A, et al. Climate change and habitat fragmentation drive the occurrence of Borrelia burgdorferi, the agent of Lyme disease, at the northeastern limit of its distribution. Evol Appl. 2014;7:750–64.

11. Keirans JE, Litwak TR. Pictorial Key to the Adults of Hard Ticks, Family Ixodidae (Ixodida: Ixodoidea), East of the Mississippi River. J Med Entomol. 1989;26:435–48.

12. Cooley RA, Kohls GM. The genus Amblyomma (Ixodidae) in the United States. J Parasitol [Internet]. 1944;30. Available from: https://doi.org/10.2307/3272571

13. Keirans JE, Clifford CM. The Genus Ixodes in the United States: A Scanning Electron Microscope Study and Key to the Adults. J Med Entomol. 1978;15:1–38.

14. Yunker CE, Keirans JE, Clifford CM, Easton EA. Dermacentor ticks (Acari: Ixodoidea: Ixodidae) of the New World: a scanning electron microscope atlas. Proc Entomol Soc Wash. 1986;88:609–27.

15. Durden LA, Keirans JE. Description of the larva, diagnosis of the nymph and female based on scanning electron microscopy, hosts, and distribution of Ixodes (Ixodes) venezuelensis. Med Vet Entomol. 1994;8:310–6.

16. Durden LA, Keirans JE. Nymphs of the genus Ixodes of the United States. Lanham, Maryland, USA: Entomological Society of America; 1996.

17. Keirans JE, Durden LA. Illustrated Key to Nymphs of the Tick Genus Amblyomma (Acari: Ixodidae) Found in the United States. J Med Entomol. 1998;35:489–95.

18. Gu Z, Gu L, Eils R, Schlesner M, Brors B. circlize Implements and enhances circular visualization in R. Bioinforma Oxf Engl. 2014;30:2811–2.

19. McLeod AI. MannKendall. 2011.

20. Anderson JF, Duray PH, Magnarelli LA. Borrelia burgdorferi and Ixodes dammini prevalent in the greater Philadelphia area. J Infect Dis. 1990;161:811–2.

21. Baer-Lehman ML, Light T, Fuller NW, Barry-Landis KD, Kindlin CM, Stewart RL. Evidence for competition between Ixodes scapularis and Dermacentor albipictus feeding concurrently on white-tailed deer. Exp Appl Acarol. 2012;58:301–14.

22. Brown SM, Lehman PM, Kern RA, Henning JD. Detection of Borrelia burgdorferi and Anaplasma phagocytophilum in the black-legged tick, Ixodes scapularis, within southwestern Pennsylvania. J Vector Ecol J Soc Vector Ecol. 2015;40:180–3.

23. Campagnolo ER, Tewari D, Farone TS, Livengood JL, Mason KL. Evidence of Powassan/deer tick virus in adult black-legged ticks (Ixodes scapularis) recovered from hunter-harvested white-tailed deer (Odocoileus virginianus) in Pennsylvania: A public health perspective. Zoonoses Public Health. 2018;65:589–94.

24. Courtney JW, Dryden RL, Montgomery J, Schneider BS, Smith G, Massung RF. Molecular characterization of Anaplasma phagocytophilum and Borrelia burgdorferi in Ixodes scapularis ticks from Pennsylvania. J Clin Microbiol. 2003;41:1569–73.

25. Crowder CD, Carolan HE, Rounds MA, Honig V, Mothes B, Haag H, et al. Prevalence of Borrelia miyamotoi in Ixodes ticks in Europe and the United States. Emerg Infect Dis. 2014;20:1678–82.

26. Daniels TJ, Fish D, Levine JF, Greco MA, Eaton AT, Padgett PJ, et al. Canine exposure to Borrelia burgdorferi and prevalence of Ixodes dammini (Acari: Ixodidae) on deer as a measure of Lyme disease risk in the northeastern United States. J Med Entomol. 1993;30:171–8.

27. Devevey G, Brisson D. The effect of spatial heterogenity on the aggregation of ticks on white-footed mice. Parasitology. 2012;139:915–25.

28. Dick CW, Gannon MR, Little WE, Patrick MJ. Ectoparasite associations of bats from central Pennsylvania. J Med Entomol. 2003;40:813–9.

29. Edwards MJ, Barbalato LA, Makkapati A, Pham KD, Bugbee LM. Relatively low prevalence of Babesia microti and Anaplasma phagocytophilum in Ixodes scapularis ticks collected in the Lehigh Valley region of eastern Pennsylvania. Ticks Tick-Borne Dis. 2015;6:812–9.

30. Han GS, Stromdahl EY, Wong D, Weltman AC. Exposure to Borrelia burgdorferi and other tick-borne pathogens in Gettysburg National Military Park, South-Central Pennsylvania, 2009. Vector Borne Zoonotic Dis Larchmt N. 2014;14:227–33.

31. Hutchinson ML, Strohecker MD, Simmons TW, Kyle AD, Helwig MW. Prevalence Rates of Borrelia burgdorferi (Spirochaetales: Spirochaetaceae), Anaplasma phagocytophilum (Rickettsiales: Anaplasmataceae), and Babesia microti (Piroplasmida: Babesiidae) in Host-Seeking Ixodes scapularis (Acari: Ixodidae) from Pennsylvania. J Med Entomol. 2015;52:693–8.

32. Lo Re V, Occi JL, MacGregor RR. Identifying the vector of Lyme disease. Am Fam Physician.2004;69:1935–7.

33. Lord RD, Lord VR, Humphreys JG, McLean RG. Distribution of Borrelia burgdorferi in host mice in Pennsylvania. J Clin Microbiol. 1994;32:2501–4.

34. Magnarelli LA, Stafford KC, Mather TN, Yeh MT, Horn KD, Dumler JS. Hemocytic rickettsia-like organisms in ticks: serologic reactivity with antisera to Ehrlichiae and detection of DNA of agent of human granulocytic ehrlichiosis by PCR. J Clin Microbiol. 1995;33:2710–4.

35. Miller MJ, Esser HJ, Loaiza JR, Herre EA, Aguilar C, Quintero D, et al. Molecular Ecological Insights into Neotropical Bird-Tick Interactions. PloS One. 2016;11:e0155989.

36. Rogers MB, Cui L, Fitch A, Popov V, Travassos da Rosa APA, Vasilakis N, et al. Whole genome analysis of sierra nevada virus, a novel mononegavirus in the family nyamiviridae. Am J Trop Med Hyg. 2014;91:159–64.

37. Sakamoto JM, Ng TFF, Suzuki Y, Tsujimoto H, Deng X, Delwart E, et al. Bunyaviruses are common in male and female Ixodes scapularis ticks in central Pennsylvania. PeerJ. 2016;4:e2324.

38. Sakamoto JM, Goddard J, Rasgon JL. Population and demographic structure of Ixodes scapularis Say in the eastern United States. PloS One. 2014;9:e101389.

39. Schoelkopf L, Hutchinson CE, Bendele KG, Goff WL, Willette M, Rasmussen JM, et al. New ruminant hosts and wider geographic range identified for Babesia odocoilei (Emerson and Wright 1970). J Wildl Dis. 2005;41:683–90.

40. Serfass TL, Rymon LM, Brooks RP. Ectoparasites from river otters in Pennsylvania. J Wildl Dis. 1992;28:138–40.

41. Shock BC, Moncayo A, Cohen S, Mitchell EA, Williamson PC, Lopez G, et al. Diversity of piroplasms detected in blood-fed and questing ticks from several states in the United States. Ticks Tick-Borne Dis. 2014;5:373–80.

42. Simmons TW, Shea J, Myers-Claypole MA, Kruise R, Hutchinson ML. Seasonal Activity, Density, and Collection Efficiency of the Blacklegged Tick (Ixodes scapularis) (Acari: Ixodidae) in Mid-Western Pennsylvania. J Med Entomol. 2015;52:1260–9.

43. Snetsinger R, Jacobs SB, Kim KC, Tavris D. Extension of the range of Dermacentor variabilis (Acari: Ixodidae) in Pennsylvania. J Med Entomol. 1993;30:795–8.

44. Springer YP, Jarnevich CS, Barnett DT, Monaghan AJ, Eisen RJ. Modeling the Present and Future Geographic Distribution of the Lone Star Tick, Amblyomma americanum (Ixodida: Ixodidae), in the Continental United States. Am J Trop Med Hyg. 2015;93:875–90.

45. Steiner FE, Pinger RR, Vann CN, Grindle N, Civitello D, Clay K, et al. Infection and co-infection rates of Anaplasma phagocytophilum variants, Babesia spp., Borrelia burgdorferi, and the rickettsial endosymbiont in Ixodes scapularis (Acari: Ixodidae) from sites in Indiana, Maine, Pennsylvania, and Wisconsin. J Med Entomol. 2008;45:289–97.

46. Stromdahl E, Hamer S, Jenkins S, Sloan L, Williamson P, Foster E, et al. Comparison of phenology and pathogen prevalence, including infection with the Ehrlichia muris-like (EML) agent, of Ixodes scapularis removed from soldiers in the midwestern and the northeastern United States over a 15 year period (1997-2012). Parasit Vectors. 2014;7:553.

47. Waits K, Edwards MJ, Cobb IN, Fontenele RS, Varsani A. Identification of an anellovirus and genomoviruses in ixodid ticks. Virus Genes. 2018;54:155–9.

48. Yeh MT, Bak JM, Hu R, Nicholson MC, Kelly C, Mather TN. Determining the duration of Ixodes scapularis (Acari: Ixodidae) attachment to tick-bite victims. J Med Entomol. 1995;32:853–8.

49. Snetsinger RJ. Distribution of Ticks and Tick-Borne Diseases in Pennsylvania. University Park, PAJ: Pennsylvania State University, College of Agriculture, Agricultural Experiment Station,; 1968.

50. Chen X, Xu S, Yu Z, Guo L, Yang S, Liu L, et al. Multiple lines of evidence on the genetic relatedness of the parthenogenetic and bisexual Haemaphysalis longicornis (Acari: Ixodidae). Infect Genet Evol J Mol Epidemiol Evol Genet Infect Dis. 2014;21:308–14.

51. Chinuki Y, Ishiwata K, Yamaji K, Takahashi H, Morita E. Haemaphysalis longicornis tick bites are a possible cause of red meat allergy in Japan. Allergy. 2016;71:421–5.

52. Rainey T, Occi JL, Robbins RG, Egizi A. Discovery of Haemaphysalis longicornis (Ixodida: Ixodidae) Parasitizing a Sheep in New Jersey, United States. J Med Entomol. 2018;55:757–9.

53. Bishopp FC, Trembley HL. Distribution and Hosts of Certain North American Ticks. J Parasitol. 1945;31:1–54.

54. Pretzman C, Daugherty N, Poetter K, Ralph D. The distribution and dynamics of Rickettsia in the tick population of Ohio. Ann N Y Acad Sci. 1990;590:227–36.

55. Wang P, Glowacki MN, Hoet AE, Needham GR, Smith KA, Gary RE, et al. Emergence of Ixodes scapularis and Borrelia burgdorferi, the Lyme disease vector and agent, in Ohio. Front Cell Infect Microbiol [Internet]. 2014 [cited 2017 Nov 19];4. Available from: https://www.frontiersin.org/articles/10.3389/fcimb.2014.00070/full#h4

56. Brites-Neto J, Duarte KMR, Martins TF. Tick-borne infections in human and animal population worldwide. Vet World. 2015;8:301–15.

57. ProMED-mail. Rocky mountain spotted fever - USA (03): (PA) fatal [Internet]. 2014 Sep. Available from: https://www.promedmail.org/post/2757657

58. Levin ML, Zemtsova GE, Killmaster LF, Snellgrove A, Schumacher LBM. Vector competence of Amblyomma americanum (Acari: Ixodidae) for Rickettsia rickettsii. Ticks Tick-Borne Dis. 2017;8:615–22.

59. Savage HM, Godsey MS, Panella NA, Burkhalter KL, Manford J, Trevino-Garrison IC, et al. Surveillance for Tick-Borne Viruses Near the Location of a Fatal Human Case of Bourbon Virus (Family Orthomyxoviridae: Genus Thogotovirus) in Eastern Kansas, 2015. J Med Entomol. 2018;

60. Commins SP, Platts-Mills TAE. Tick bites and red meat allergy. Curr Opin Allergy Clin Immunol. 2013;13:354–9.

61. Guglielmone AA, Robbins RG, Apanaskevich DA, Petney TN, Estrada-Peña A, Horak IG. The Hard Ticks of the World [Internet]. Dordrecht: Springer Netherlands; 2014 [cited 2018 Mar 14]. Available from: http://link.springer.com/10.1007/978-94-007-7497-1

62. Parker RR, Pickens EG, Lackman DB, Bell EJ, Thraikill FB. Isolation and Characterization of Rocky Mountain Spotted Fever Rickettsiae from the Rabbit Tick Haemaphysalis leporis-palustris Packard. Public Health Rep 1896-1970. 1951;66:455–63.

63. Gabriele-Rivet V, Arsenault J, Badcock J, Cheng A, Edsall J, Goltz J, et al. Different Ecological Niches for Ticks of Public Health Significance in Canada. PLOS ONE. 2015;10:e0131282.

64. Schneider SC, Parker CM, Miller JR, Page Fredericks L, Allan BF. Assessing the Contribution of Songbirds to the Movement of Ticks and Borrelia burgdorferi in the Midwestern United States During Fall Migration. EcoHealth. 2015;12:164–73.

65. Burg JG. Seasonal activity and spatial distribution of host-seeking adults of the tick Dermacentor variabilis. Med Vet Entomol. 2001;15:413–21.

66. Kollars TM, Oliver JH, Durden LA, Kollars PG. Host Associations and Seasonal Activity of Amblyomma americanum (Acari: Ixodidae) in Missouri. J Parasitol. 2000;86:1156–9.

67. Ko RC. Biology of Ixodes cookei Packard (Ixodidae) of groundhogs (Marmota monax Erxleben). Can J Zool. 1972;50:433–6.

68. McLean DM, Best JM, Mahalingam S, Chernesky MA, Wilson WE. Powassan Virus: Summer Infection Cycle, 1964. 1965.

69. Ryder JW, Pinger RR, Glancy T. Inability of Ixodes cookei and Amblyomma americanum nymphs (Acari: Ixodidae) to transmit Borrelia burgdorferi. J Med Entomol. 1992;29:525–30.

70. Brillhart DB, Fox LB, Upton SJ. Ticks (Acari: Ixodidae) collected from small and medium-sized Kansas mammals. J Med Entomol. 1994;31:500–4.

71. Cohen SB, Freye JD, Dunlap BG, Dunn JR, Jones TF, Moncayo AC. Host associations of Dermacentor, Amblyomma, and Ixodes (Acari: Ixodidae) ticks in Tennessee. J Med Entomol. 2010;47:415–20.

72. Merten HA, Durden LA. A state-by-state survey of ticks recorded from humans in the United States. J Vector Ecol J Soc Vector Ecol. 2000;25:102–13.

73. Barker IK, Lindsay LR, Campbell GD, Surgeoner GA, McEwen SA. The groundhog tick Ixodes cookei (Acari: Ixodidae): a poor potential vector of Lyme borreliosis. J Wildl Dis. 1993;29:416–22.

74. Wright CL, Sonenshine DE, Gaff HD, Hynes WL. Rickettsia parkeri Transmission to Amblyomma americanum by Cofeeding with Amblyomma maculatum (Acari: Ixodidae) and Potential for Spillover. J Med Entomol. 2015;52:1090–5.

75. Main AJ, Carey AB, Downs WG. POWASSAN VIRUS IN Ixodes cookei AND MUSTELIDAE IN NEW ENGLAND. J Wildl Dis. 1979;15:585–91.

76. Eisen RJ, Eisen L, Beard CB. County-Scale Distribution of Ixodes scapularis and Ixodes pacificus (Acari: Ixodidae) in the Continental United States. J Med Entomol. 2016;53:349–86.

77. Krawczak FS, Nieri-Bastos FA, Nunes FP, Soares JF, Moraes-Filho J, Labruna MB. Rickettsial infection in Amblyomma cajennense ticks and capybaras (Hydrochoerus hydrochaeris) in a Brazilian spotted fever-endemic area. Parasit Vectors [Internet]. 2014;7. Available from: https://doi.org/10.1186/1756-3305-7-7

78. Kohls GM, Hoogstraal H, Clifford CM, Kaiser MN. The Subgenus Persicargas (Ixodoidea, Argasidae, Argas). 9. Redescription and New World Records of Argas (P.) persicus (Oken), and Resurrection, Redescription, and Records of A. (P.) radiatus Railliet, A. (P.) sanchezi Duges, and A. (P.) miniatus Koch, New World Ticks Misidentified as A. (P.) persicus. Ann Entomol Soc Am. 1970;63:590–606.

79. Keirans JE, Durden LA. Invasion: Exotic Ticks (Acari: Argasidae, Ixodidae) Imported into the United States. A Review and New Records. J Med Entomol. 2001;38:850–61.

80. Muñoz-Leal S, Venzal JM, Nava S, Reyes M, Martins TF, Leite RC, et al. The geographic distribution of Argas (Persicargas) miniatus and Argas (Persicargas) persicus (Acari: Argasidae) in America, with morphological and molecular diagnoses from Brazil, Chile and Cuba. Ticks Tick-Borne Dis. 2018;9:44–56.

81. Kohls GM, Hoogstraal H. Observations on the Subgenus Argas (Ixodoidea, Argasidae, Argas) 2. A. Cooleyi, New Species, from Western North American Birds1. Ann Entomol Soc Am. 1960;53:625–31.

82. Cortinas R, Spomer SM. Occurrence and County-Level Distribution of Ticks (Acari: Ixodoidea) in Nebraska using Passive Surveillance. J Med Entomol. 2014;51:352–9.

83. Barrett AW, Noden BH, Gruntmeir JM, Holland T, Mitcham JR, Martin JE, et al. County Scale Distribution of Amblyomma americanum (Ixodida: Ixodidae) in Oklahoma: Addressing Local Deficits in Tick Maps Based on Passive Reporting. J Med Entomol. 2015;52:269–73.

84. Xu G, Mather TN, Hollingsworth CS, Rich SM. Passive Surveillance of Ixodes scapularis (Say), Their Biting Activity, and Associated Pathogens in Massachusetts. Vector Borne Zoonotic Dis. 2016;16:520–7.

85. Springer YP, Eisen L, Beati L, James AM, Eisen RJ. Spatial Distribution of Counties in the Continental United States With Records of Occurrence of Amblyomma americanum (Ixodida: Ixodidae). J Med Entomol. 2014;51:342–51.

